# Uncovering the latent structure of interwoven population and temporal codes

**DOI:** 10.64898/2026.05.11.724260

**Authors:** Zachary Friedenberger, Yiwei Cao, Richard Naud

## Abstract

Population analysis methods have become standard for navigating the complexity of neural data. However, these methods often assume a rate code, neglecting information encoded in the precise timing of spikes. Critically, additional information encoded in bursts of action potentials may be missed. Here, we develop a probabilistic latent-variable framework that disentangles the factors associated with bursts and individual spikes. This enables burst codes to be investigated directly from the structure of the data, without requiring external covariates. We demonstrate that analyzing firing rates alone obscures the latent structure and factors underlying bursts. Applying our method to simulated and experimental data, we show that it can infer the correct latent structure and be used to test for the presence of burst coding. By merging the population and burst coding perspectives, we provide a framework for linking changes in bursting to internal variables involved in attention, perception, and learning.

## 1 Introduction

Understanding how to read the neural code is crucial to our interpretation of how the brain operates [1–5]. As modern technologies now enable the simultaneous recording of large populations of neurons, *population coding* has emerged as a leading candidate [6–8]. In this code, information is encoded in the coordinated activity of many individual neurons and can often be compactly described using a small set of latent factors [9–12]. Therefore, dimensionality reduction methods have become essential for making sense of neural data and inferring the factors relevant to sensory processing and behaviour [13–16].

Underlying these methods is the assumption of a *rate code*, in which single neurons contribute to the population only through their firing rate. While this has been fruitful in relating neural activity to behaviour, it necessarily glosses over temporal spike patterns such as bursts of action potentials, which are known to strongly influence synaptic dynamics and plasticity [5, 17–20]. Critically, factors associated with bursts may be overlooked, a potentially serious omission as bursts are believed to play a role in perception, attention, and learning [5, 21–27]. Solely focusing on firing rates may therefore miss additional factors, providing an incomplete view of the neural code. However, inferring these additional factors and disentangling them from those associated with firing rates requires new principled methods.

To fill this gap, we introduce *Multiplexed Factor Analysis* (MFA), a probabilistic latent-variable modelling framework for analyzing interwoven population and temporal codes. MFA decomposes spike trains into bursts and events (i.e., individual spikes and bursts), enabling the factors underlying burstiness and event rate to be disentangled. By treating bursts as a unique syllable of the neural code with their own factors, we address a fundamental limitation of previous work, moving beyond descriptive modelling of spiking statistics and towards biological interpretability [28]. This is an important step for testing theories of credit assignment and information routing that rely on such codes [29, 30].

## 2 Background

### 2.1 Factor analysis for neural population activity

Factor Analysis (FA) is a linear-Gaussian latent-variable model whose goal is to explain the data covariance structure via a decomposition into shared and private components [31]. FA is typically applied to neural activity in the form of spike counts, *a*, observed across a population of *N* neurons [14, 32, 33]. As neurons have Poissonlike statistics *in vivo*, spike counts are variance-stabilized using the square-root transform, 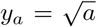, to approximate a Gaussian [14, 34]. The standard FA model is,

,
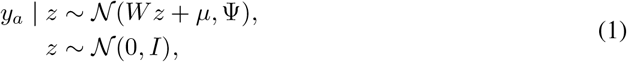

where *y*_*a*_ ∈ ℝ^*N*^ are the transformed spike counts, *z* ∈ ℝ^*K*^ are the factors, *W* ∈ ℝ^*N* ×*K*^ are the factor loading weights whose columns define a low-dimensional subspace capturing the covariability between neurons, *µ* ∈ ℝ^*N*^ defines the mean of each neuron, and Ψ is a diagonal matrix capturing the private variability of each neuron.

FA and its extensions yield low-dimensional representations of firing rates, in which factors describe shared fluctuations, and their loadings define the population structure. Importantly, these factors are inferred directly from neural activity without detailed knowledge of what is being encoded, enabling exploratory analyses at the expense of complex inference (see Section 4.1 for inference details).

### 2.2 Analyzing bursts of action potentials in single neurons

Bursts of action potentials, defined as a sudden increase in the firing frequency followed by a period of silence, often display an alteration in spike waveform and a prolonged depolarization of the subthreshold membrane potential [5]. Prior work has measured the burstiness of spike trains using several different methods [23, 26, 35–37]. However, to interpret burstiness, these methods rely on external covariates to determine when to look for fluctuations in burstiness and to understand their relevance, an approach complicated by the heterogeneity of single-neuron responses observed *in vivo* and by the possibility that bursts encode internal variables.

In one method, spike times are classified into bursts and events based on inter-spike intervals (ISIs), and burstiness is measured using a burst fraction, defined as the ratio of bursts to events [26, 29]. This distinction is motivated by the fact that pyramidal neurons in the cortex receive segregated inputs to their somatic and apical dendrites, with the simultaneous occurrence of both triggering distinctive bursts [38–40]. Importantly, theoretical work has shown that for this conjunctive mechanism, somatic and dendritic inputs are expected to correlate with event rate and burst fraction, respectively [29]. This enables neural populations to *multiplex* two distinct streams of information in their spikes. Although the distributions of event rate and burst fraction across neural populations have been investigated [26, 27], these analyses relied heavily on single-neuron response classifications and population averaging, and did not consider the latent structure underlying bursts.

An additional challenge when investigating burst codes is that fluctuations in burstiness may be explained by fluctuations in firing rate.^1^ In this case, some of the classified bursts are simply spikes that occur close in time and are not expected to encode additional information. Therefore, determining whether fluctuations in burstiness arise independently of firing rate requires careful comparisons with null models. While prior work has considered an inhomogeneous Poisson null model [26], a general approach is needed to distinguish between bursts driven by fluctuations in firing rate and those driven by independent factors.

## 3 Multiplexed factor analysis

To address the shortcomings of prior analysis methods, we developed a probabilistic latent-variable framework for modelling the statistics of bursts and events, which leverages the shared variability across neurons to 1) uncover latent factors and 2) test against a null model. We refer to our framework as *Multiplexed Factor Analysis* (MFA), as it is designed to disentangle and recover multiplexed factors from neural population data. We considered two model variants within the MFA framework. The first, which we refer to as the *base model*, is distinct from the null model, whereas the second, which we refer to as the *hybrid model*, contains the null model as a special case. While not all bursts are generated via a conjunctive mechanism or require dendrites [5], we use it to motivate the emission distribution of our model.^2^

For application to spike time data, we defined bursts as two or more spikes with an ISI less than 15 ms [26, 29], designating the time of the first spike as the time of a burst (Fig. 1a left). Events were defined as the set of all singlet spikes and bursts.

**Figure 1:**
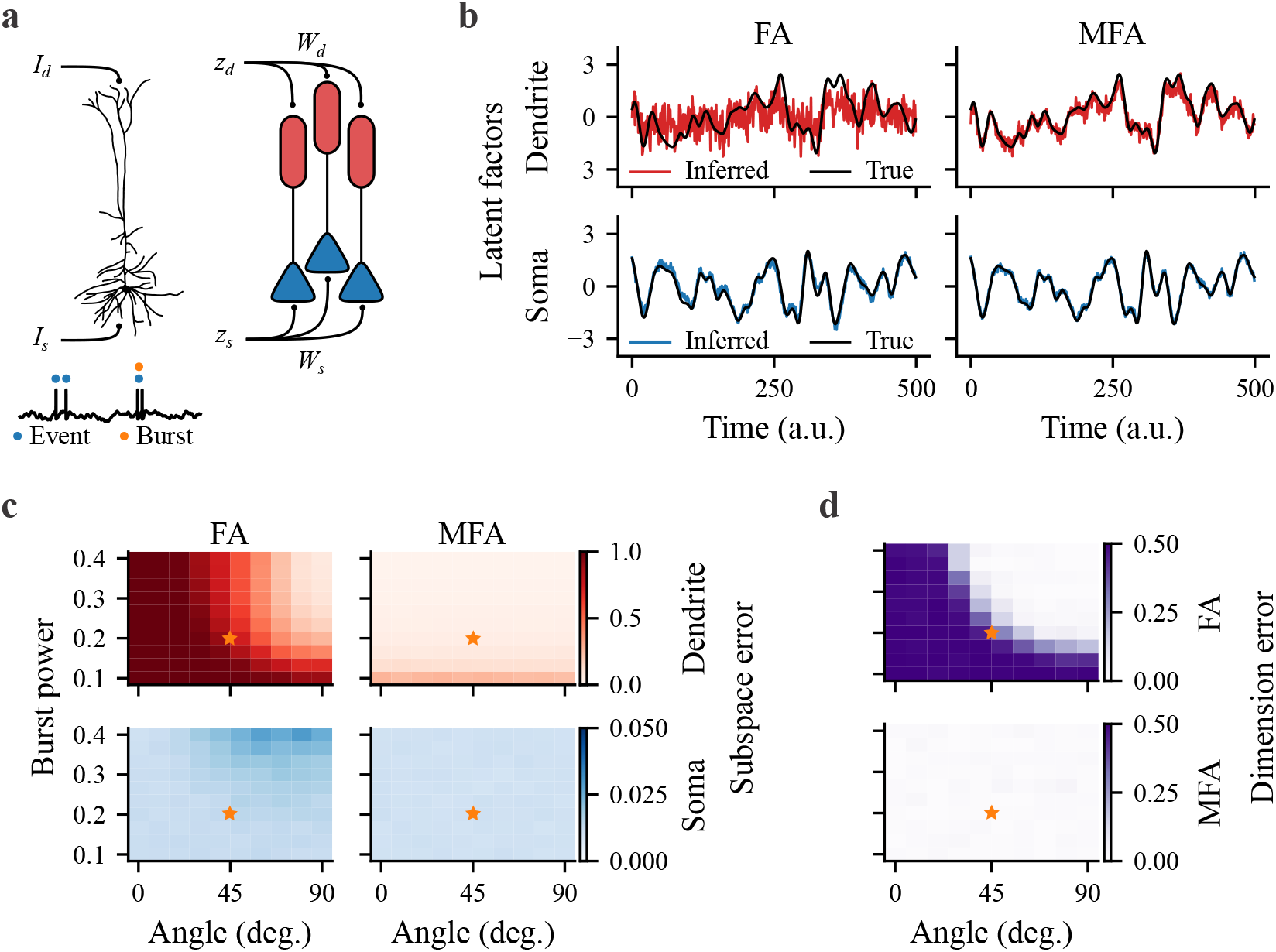
MFA reliably recovers multiplexed factors from simulated data. **a**, Thick-tufted pyramidal neuron receiving somatic *I*_*s*_ and apical dendritic inputs *I*_*d*_ (left). Spike times classified into events (blue dots) and bursts (orange dots). Illustration of the MFA base model as a population of neurons driven by somatic *z*_*s*_ and dendritic *z*_*d*_ factors through the loading weights *W*_*s*_ and *W*_*d*_, respectively (right). **b**, The factors inferred from the simulated data. FA on all spike counts (left), and MFA on event and burst counts (right). **c-d**, Subspace and dimension error as a function of the angle between subspaces and burst power. Orange stars denote the parameters used in **b**.

### 3.1 Base model

We considered a model for the number of events and bursts observed from a population of *N* neurons (Fig. 1a, right), where each event is randomly converted into a burst with an underlying burst probability. Neurons are conditionally independent given an event rate *λ*_*e*_ and burst probability *ρ*,

,
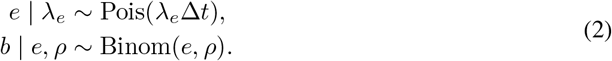

The number of events *e*, in a time bin of size Δ*t*, is Poisson distributed with event-rate *λ*_*e*_, and the number of bursts *b* is distributed according to a Binomial distribution with *e* trials and burst probability *ρ*. The number of events and bursts is related to the total spike count, *a*, by the equation *a* = *e* + (*n* − 1)*b*, where *n* is the number of spikes per burst (burst length), and the (*n* − 1) term accounts for the subsequent spikes in a burst that are not classified as events.

Motivated by the observation that event rate and burst probability can be independently modulated by somatic and dendritic inputs, respectively [29], we modelled each neuron’s event rate and burst probability as a combination of independent factors,

,
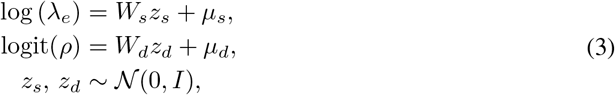

where 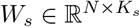 and 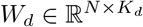 are the factor loading weights, 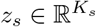 and 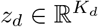 are the factors, *µ*_*s*_, 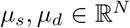 define the means, and nonlinear functions are applied point-wise to map each linear-Gaussian model onto the correct domain. The loadings define *K*_*s*_ and *K*_*d*_ dimensional linear subspaces for event rate and burst probability, which we refer to as the somatic and dendritic subspaces. Additionally, we refer to *z*_*s*_ and *z*_*d*_ as somatic and dendritic factors. However, the factors *z*_*d*_ and their loadings *W*_*d*_ may also capture nondendritic influences on burst probability.

### 3.2 Null model

To determine whether the observed latent structure in bursting could be explained by event rates alone, we considered a null model,

,
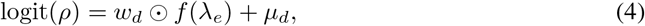

where neuron-specific dendritic weights *w*_*d*_ ∈ ℝ^*N*^ scale a function of the event rate point-wise. The resulting model is highly constrained, as fluctuations in burst probability can be predicted only from fluctuations in event rate. The model therefore generalizes the inhomogeneous Poisson null model by additionally accounting for population structure. Here, we use the centred log event rate, 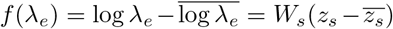, where the bar is the sample mean. Somatic loadings *W*_*s*_ and factors *z*_*s*_, however, are inferred using events only and are fixed when inferring the neuron-specific *w*_*d*_ using bursts. In our analysis, we also considered the trivial case in which *w*_*d*_ is set to zero, corresponding to neurons with a constant burst probability.

### 3.3 Hybrid model

As neurons receiving coincident somatic and dendritic inputs may also produce occasional short bursts due to fluctuations in event rate, and not all neurons in a population may employ burst coding, we considered a hybrid model that incorporated both rate-dependent and independent factor components,

,
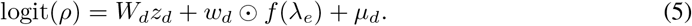

Where burst probability depends on both the dendritic factors *z*_*d*_ and the event rate *λ*_*e*_. Importantly, the hybrid model nests both the base and null models, recovering the former when *w*_*d*_ = 0 and the latter when *K*_*d*_ = 0. This nested structure allows us to determine whether dendritic factors improve predictive performance beyond what can be explained by fluctuations in event rate.

### 3.4 Relationship to prior models

#### Burst generation models

The emission model in Eq. 2 has been implicitly assumed in studies of conjunctive burst codes [26, 27]. Its maximum likelihood estimators for event rate and burst probability are the trial-averaged event rate and burst fraction, respectively (see Appendix A). However, burst fractions are ill-defined for sparse activity, obscure trial-to-trial variability, and do not capture population structure without external covariates. MFA addresses these limitations by inferring population structure and burst probabilities without explicitly computing burst fractions or requiring external covariates. Alternatively, burst generation can be described by marked-point processes [29, 41]. The emission model used in MFA can be viewed as a coarse-grained marked-point process that neglects spike-history effects, which we leave as an extension for future work (see Discussion).

#### Factor analysis and latent variable models

The event and burst components of MFA can be recognized as Poisson and Binomial FA models, respectively. Latent variable models with Poisson and exponential family emission distributions have been used extensively to model firing rates [11, 42–46]. However, these models primarily focused on dynamic latent variables. Here, we consider only the static version and leave the dynamic extension for future work (see Discussion). Overall, each MFA model can be seen as a multi-view FA model [47], where events and bursts are treated as separate views of the spike-generation process with a specific conditional structure.

#### Gain-modulation models

Closest to MFA are models that decompose firing rates into a product between a stimulus-driven component and a multiplicative gain component that varies over time. Prior models can be separated into those that focused on single-neuron responses [28, 48] and those that investigated shared gain across neural populations [16, 32, 49]. As modulations in burst probability can also be viewed as a multiplicative gain on the event rate, we expect these models to yield results similar to ours in some cases. However, MFA is defined by a distinct generative model that provides a biologically interpretable vocabulary for investigating gain modulation.

## 4 Methods

### 4.1 Model inference

For each dataset, we inferred the somatic and dendritic subspaces using two different approaches depending on the statistics of the data. For the first approach, we approximated the emission distributions in Eq. 2 as Gaussians to use the expectation-maximization (EM) algorithm [14]. This procedure is fast but requires explicit computation of burst fractions and thus performs poorly when events are sparse. For the second approach, we considered variational inference (VI) [50–52]. While slower, VI does not require computing burst fractions, performs better when events are sparse, and offers greater flexibility for quickly testing multiple models.

#### Gaussian-approximate expectation maximization

Event and burst counts were used to construct transformed variables using the square-root, 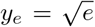, and arcsine-square-root transforms, 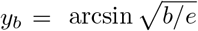 [34]. However, because the per-sample burst fraction, *b/e*, is undefined when *e* = 0, we set *b/e* = 0 for *e* = 0. The transformed variables were then fit to separate Gaussian FA models (Eq. 1). Since the posterior of Eq. 1 can be computed analytically, the EM algorithm was used to infer the subspaces [31]. This approach performs poorly for sparse events because a large proportion of samples have undefined burst fractions, leading to *y*_*b*_ being set to zero in an ad hoc manner (see Fig. S6 for a comparison between approaches).

#### Full-model variational inference

Since the posterior of the full model is intractable, approximate inference techniques are required to maximize the marginal log-likelihood, ln *p*(*D*|*θ*), where *θ* = {*W*_*s*_, *W*_*d*_, *w*_*d*_, *µ*_*s*_, *µ*_*d*_} are the model parameters and *D* is the dataset, 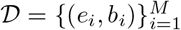. Here, we used a black-box VI approach with a mean-field assumption on the variational distribution [53].

VI introduces an approximate posterior distribution *q*(*z* | *ϕ*) for the latent variables *z*, with variational parameters *ϕ*. The model parameters *θ* and the variational parameters *ϕ* are inferred jointly by optimizing the evidence lower bound (ELBO), which is a lower bound on the marginal log-likelihood, ln *p*(*x*|*θ*) ≥ ℒ (*θ, ϕ*), where *x* = (*e, b*). The ELBO is given by,

,
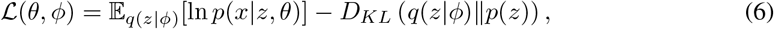

where the first term is the expected log-likelihood and the second term is the KL-divergence between the approximate posterior and the prior. We assumed that the variational posterior factorizes over the somatic and dendritic factors, *q*(*z* | *ϕ*) = *q*(*z*_*s*_ | *ϕ*_*s*_)*q*(*z*_*d*_ | *ϕ*_*d*_), with each distribution further factorized across latent dimensions. The approximate posterior of each data sample is 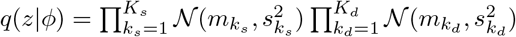, where the means and variances define the variational parameters to be optimized, 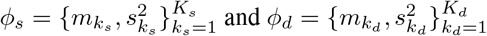.

We optimized Eq. 6, using a black-box approach where the expectation was approximated by Monte Carlo (MC) sampling from the approximate posterior and gradients were computed using auto-differentiation. We employed the standard VI tricks to reduce estimator variance [53, 54]: the reparameterization trick, analytic Gaussian KL-divergence terms, and learning rate decay. To speed up inference and improve performance, we used the factor loading weights obtained by Gaussian-approximate EM as initial guesses for optimization.

### 4.2 Model evaluation and selection

#### Predictive log-likelihood

To compare models, we used the marginal log-likelihood evaluated on held-out data (predictive log-likelihood), defining the *score* as the mean predictive log-likelihood over held-out samples.J For a latent variable model with joint likelihood, *p*(*x, z*|*θ*), the marginal likelihood is *p*(*x*|*θ*) = *p*(*x*|*z, θ*)*p*(*z*)*dz*. For the Gaussian-approximate EM approach, this quantity is analytically tractable and easy to calculate [31]. However, as this is not the case for VI, we estimated the marginal likelihood using importance sampling [28, 55],

,
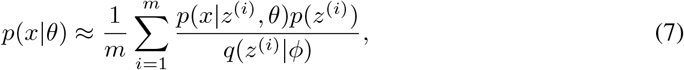

where *m* is the number of posterior samples, *z*^(*i*)^ ~ *q*(*z*|*ϕ*), and *x* = (*e, b*). In practice, the model and variational parameters are first jointly inferred from the training data. Model parameters are then fixed while new variational parameters are inferred from the test data. Equation 7 is then evaluated on the test data using the model parameters and new variational parameters.

To select the number of latent dimensions, we used K-fold cross-validation (CV) with the score relative to the simplest model as a performance metric (CV score gain). The number of dimensions was then chosen to either maximize the CV score gain or be as small as possible while retaining a high value (see Appendix E for details). Final model performance was reported as the score per event by dividing the score by the mean number of events per sample.

#### Subspace recovery metrics

For simulated data, we quantified the error between the true and estimated subspaces with a normalized projection error (subspace error). Letting *Q* and 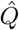 be orthonormal bases for the true *W* and estimated *Ŵ* factor loading weights, the subspace error is,

,
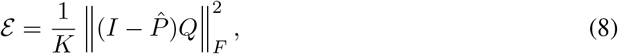

where 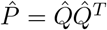 is the orthogonal projector for the estimated subspace, and ∥·∥_*F*_ is the Frobenius norm. This error is bounded such that ℰ = 0 when the subspaces coincide and ℰ = 1 when they are orthogonal. See Appendix B for a geometric interpretation and its relationship to other subspace distances. To quantify the error between the total number of true latent dimensions, *K* = *K*_*s*_ + *K*_*d*_, and the estimated number, 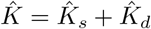, we defined the dimension error as 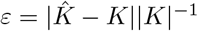.

### 4.3 Experimental datasets

To evaluate our framework on experimental data, we analyzed counts from two *in vivo* datasets. 1) the Visual Behaviour Neuropixels (VBN) dataset [56, 57], and 2) the Rodgers dataset [58]. Below, we briefly describe each dataset, with additional details provided in Appendix D.

#### VBN dataset

Mice performed a visual change-detection task while multiple Neuropixels probes simultaneously recorded activity across visual cortical areas (Fig. 3a). Recordings included sessions with familiar and novel images. We counted events and bursts during each 250 ms image presentation and pooled neurons across visual areas and cortical layers. Responses were aggregated across image identities, such that the observed variability reflects a combination of stimulus-driven and trial-to-trial fluctuations. We selected this dataset under the assumption that burstiness was more likely to be modulated during active behaviour or in response to novelty [5, 59–61].

#### Rodgers dataset

Mice performed a whisker-based shape-discrimination task, distinguishing concave from convex objects. Activity was recorded across cortical layers of the barrel cortex using a high-density silicon probe, with different barrel columns sampled across sessions (Fig. 3e). We counted events and bursts in 250 ms bins spanning ± 3 s around the choice time and primarily focused on L5b neurons as a proxy for L5 thick-tufted (L5tt) neurons, which exhibit dendritic calcium spikes and burst firing during whisker-based behaviours [22, 36, 62]. For some sessions, we pooled neurons across deep cortical layers (L5 and L6) to increase the number of simultaneously recorded neurons.

## 5 Results

We used simulations to show that MFA can uncover factors and test for the presence of burst coding by comparing against a null model. Importantly, we demonstrated that analyzing firing rates alone either misses or misinterprets these factors. Lastly, we applied MFA to the VBN and Rodgers datasets and found that a model incorporating additional dendritic factors predicted fluctuations in bursting better than the null model.

### 5.1 Factor analysis with the total number of spikes does not reliably recover dendritic factors

To compare the factors uncovered using MFA with those obtained from firing rates, we examined the recovery of one-dimensional somatic and dendritic latent factors from neural populations with conjunctive bursts (see Appendix C for simulation details). Counts were generated from the MFA base model (Eqs. 2 and 3) with random one-dimensional subspaces *W*_*s*_ and *W*_*d*_. The MFA base model was fit using events and bursts, while the FA model (Eq. 1) used the total number of spikes.

Figure 1b shows the factors inferred by the FA and MFA models. For each model, the somatic factor (blue line) closely followed the true value (black line). However, only MFA accurately recovered the dendritic factor (red line). In general, recovery of the somatic and dendritic subspaces, as measured by the subspace error (Eq. 8), heavily depends on the burst power (i.e., the standard deviation of the dendritic loading weights) and the alignment between the somatic and dendritic subspaces (Fig. 1c). Our simulation shows that MFA can recover both subspaces over a wide range of burst power and alignment angles, whereas FA recovers them only when the burst power is high and the subspaces are nearly orthogonal. Additionally, FA underestimates the correct number of latent dimensions over the same parameter range (Fig. 1d, high dimension error). This indicates that when burst coding is present, MFA can reliably infer and disentangle somatic and dendritic factors, whereas FA systematically misses or fails to fully separate them.

### 5.2 Detecting independent modulations of bursting

To demonstrate that MFA is capable of rejecting the null model in Eq. 4 and can detect the presence of burst coding, we simulated spiking activity from a population of 50 neurons containing a mixture of intact and silenced apical dendrites, reflecting the possibility that not all neurons may employ burst coding. Simulated neurons were modelled after L5tt neurons, containing a somatic and dendritic integrative compartment (Appendix C for simulation details). Conjunctive bursts were produced upon concomitant somatic and dendritic input currents. Each compartment received an input current composed of a random component shared across neurons, a constant baseline, and independent noise. The shared component defined partially aligned three-dimensional somatic and dendritic subspaces. For neurons with silenced dendrites, the coupling constant between compartments was set to zero, transforming each two-compartment model into an adaptive leaky-integrate-and-fire model.

Simulated population activity on a single trial is shown when all dendrites were intact (Fig. 2a) and silenced (Fig. 2b). When intact, bursts occurred frequently, with a mean burst fraction across the population of 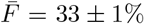 (SEM across neurons), corresponding to a regime that is optimal for multiplexing somatic and dendritic inputs in event-rate and burst fraction [29]. When silenced, the mean burst fraction decreased, as bursts were driven purely by fluctuations in somatic input, 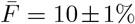. Similarly, the mean burst length also decreased (Fig. 2c), falling from 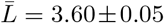 when intact, to 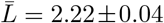 when silenced (SEM across bursts). Sweeping the fraction of neurons with intact dendrites (fraction intact) smoothly interpolated between these values (Fig. 2d).

**Figure 2:**
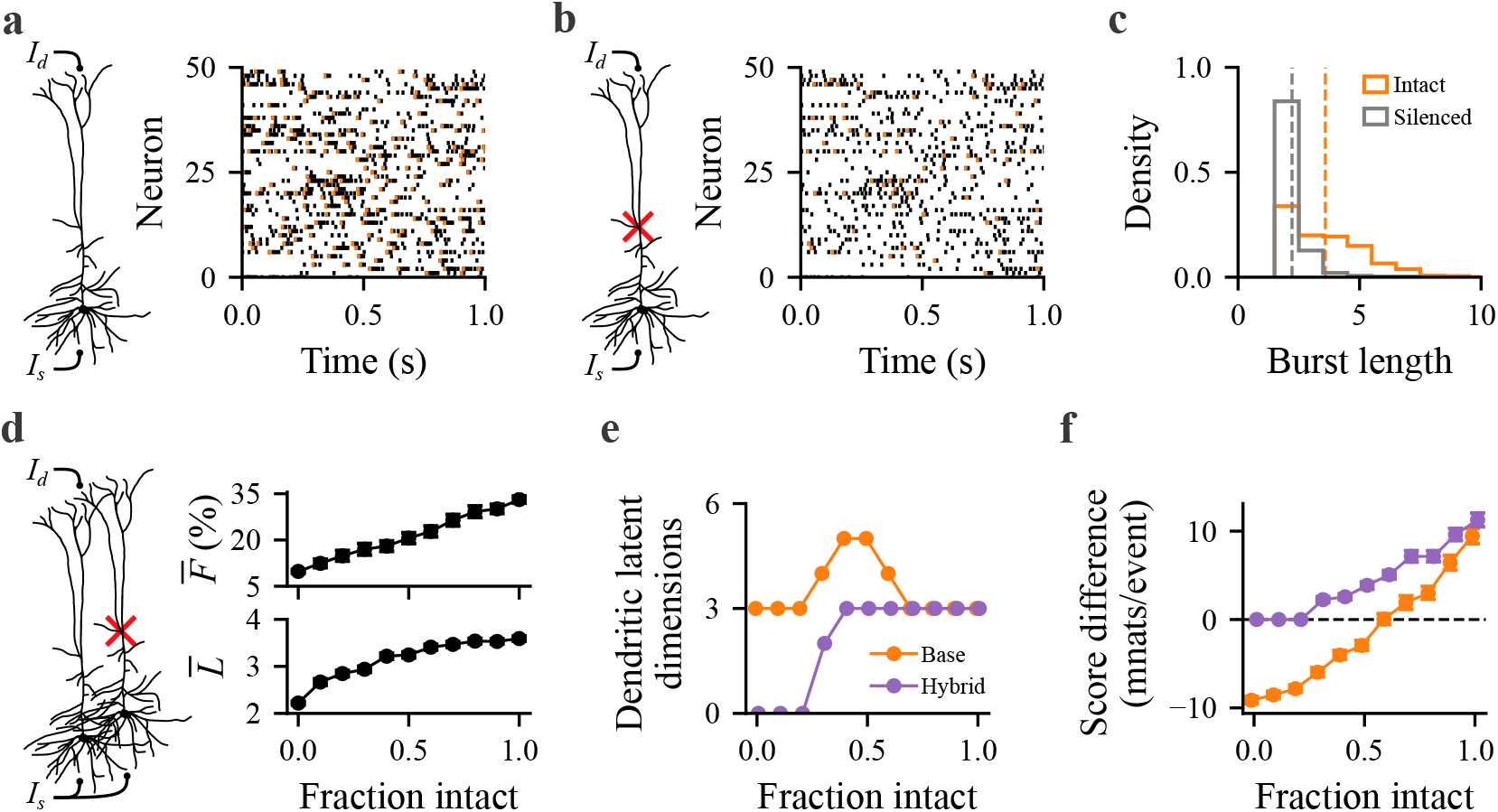
Detecting independent modulations of bursting in simulated population data. **a**, Thicktufted pyramidal neuron receiving somatic *I*_*s*_ and dendritic *I*_*d*_ input currents (left). Raster plot of the population activity on a single trial (right). Burst times are coloured orange, and all other spikes are black. One second of the trial is shown for visibility. **b**, Same as **a**, but with the dendrite silenced. **c**, Burst length distributions for intact (orange) and silenced (grey) simulations. Dashed lines indicate the mean burst length. **d**, Simulations varying the fraction of dendrites (left). The average burst fraction 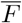 and burst length 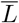 (right). **e**, The number of dendritic latent dimensions chosen for the base (orange) and hybrid (purple) models. The true dimensionality is *K*_*d*_ = 3, except when all neurons are silenced (*K*_*d*_ = 0). **f**, The difference in mean predictive log-likelihood per event (score difference). The base-null and hybrid-null model comparisons are shown in orange and purple, respectively. Error bars are 95% confidence intervals obtained via bootstrapping over test samples.

The base, hybrid, and null models were fit to each simulated dataset using counts binned over 250 ms windows. The event-rate component was fit first and then held fixed for training each burst model (10,000 training samples). CV was performed on a subset of the training data (1,000 samples) to select the number of latent dimensions for each model (see Appendix E for inference details). Models were then fit to the entire training set and evaluated on a held-out test set (2,000 test samples).

For each simulation, the event-rate component recovered the somatic subspace dimensionality (Fig. S1). When the fraction of neurons with intact dendrites was high, both the base and hybrid models recovered the correct dendritic dimensionality (Fig. 2e), and outperformed the null model (Fig. 2f, positive score difference). At intermediate fractions, however, only the hybrid model recovered the correct dimensionality and outperformed the null. The base model instead overestimated the dendritic dimensionality, requiring additional factors to approximate the rate-dependent bursting of neurons with silenced dendrites. At low fractions, the shared dendritic signal was too weak for either model to recover the dendritic subspace and outperform the null. Nevertheless, the hybrid model appropriately selected *K*_*d*_ = 0 through CV, whereas the base model continued to identify spurious factors.

These results suggest that the hybrid model provides a more reliable test for shared dendritic structure in real neural populations, while poor performance of the base model may reflect a mixture of burst-coding and non-burst-coding neurons. Importantly, they demonstrate that the null model can be rejected without prior knowledge of the specific structure or content of the burst code.

### 5.3 Application of MFA to neural data

To demonstrate that our framework could uncover factors driving bursts in real neural populations, we applied MFA to the VBN and Rodgers datasets (Section 4.3). Figure 3b shows population activity from the VBN dataset on a single trial during a novel session where the mouse correctly detected the image change. Similarly, Fig. 3f shows population activity on a single trial from the Rodgers dataset where the mouse correctly identified the presented object. The mean burst fraction across each population was 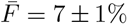 and 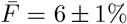 for the VBN and Rodgers data, respectively, reflecting the fact that some neurons rarely burst and bursts are sparse in time. For both sessions shown, the distribution of burst lengths was exponential-like with a mean burst length of 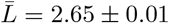 for the VBN dataset (Fig. 3c) and 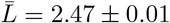 for the Rodgers dataset (Fig. 3g).

**Figure 3:**
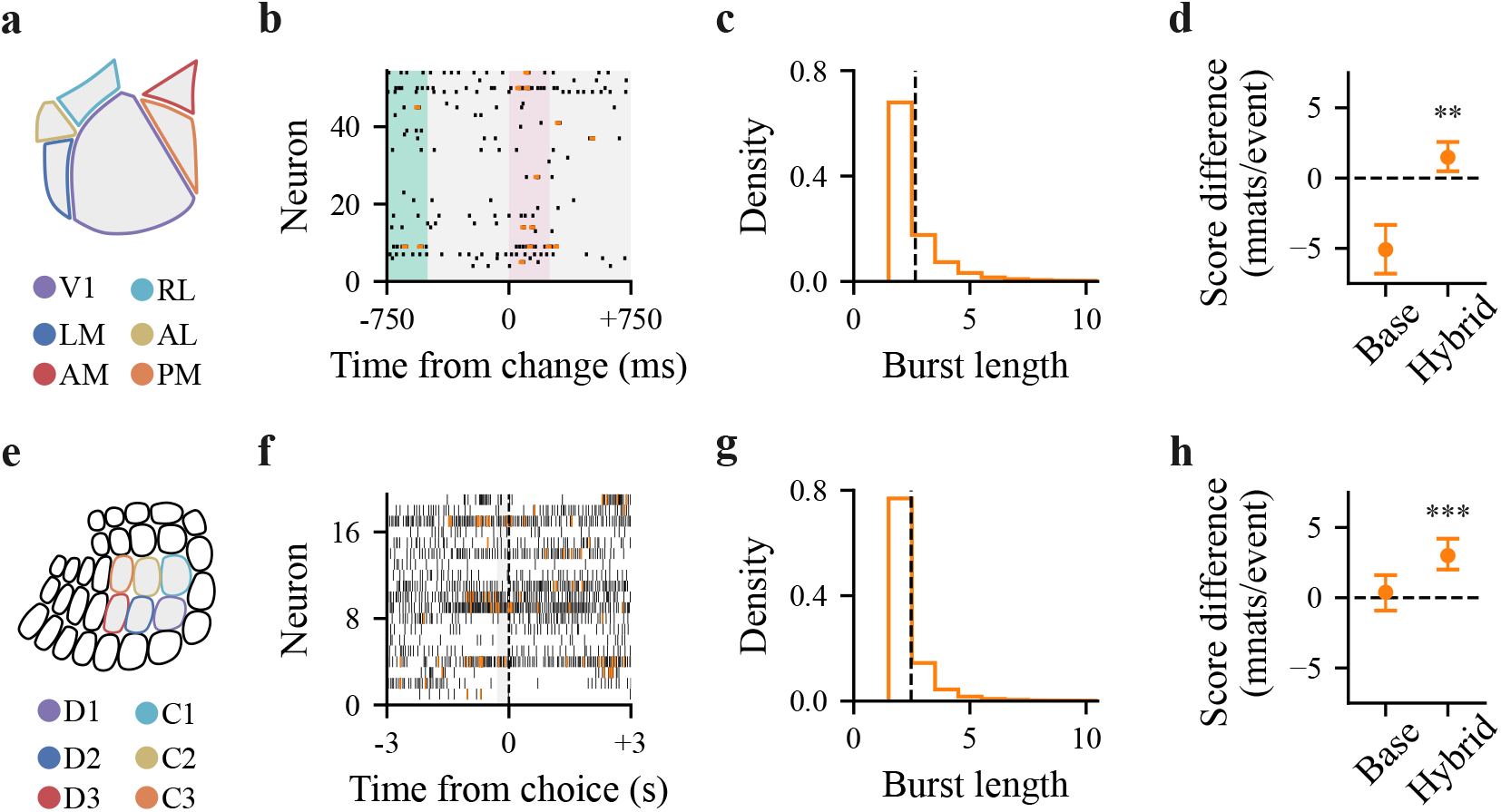
Application of MFA to neural data. **a-d**, VBN dataset. **a**, Recorded areas of mouse visual cortex. **b**, Population activity raster from a trial with correct image-change detection. Burst times are coloured orange, and all other spikes are black. Shaded regions denote the visual stimulus: pre-change image (green), change image (pink), and grey screen (grey). **c**, Distribution of burst lengths (orange), with the average burst length 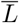 (black dashed line). **d**, The difference in mean predictive log-likelihood per event (Score difference) for base-null and hybrid-null. Error bars show the bootstrapped 95% confidence interval; ** and *** denote *p* ≤ 0.01 and *p* ≤ 0.001 for a two-sided bootstrap test of a score difference of zero. **e-h**, Rodgers dataset. **e**, Barrel columns of mouse primary somatosensory cortex. **f**, Population activity raster of L5b neurons in barrel D2 during a correct stimulus-identification trial. **g-h**, Same as **c-d** but for the Rodgers data.

The base, hybrid, and null models were fit using counts obtained from individual recording sessions (see Appendix E for inference details), split into training and test sets for evaluation (80/20 split). For brevity, we only report the full results for one session from each dataset. For the VBN dataset, we report the results from a novel session (46 neurons, 2662 training samples, and 665 test samples), while for the Rodgers dataset, we report the results for a session with activity recorded from barrel D2 where we analyzed only neurons categorized as L5b (19 neurons, 3024 training samples, and 768 test samples). Neurons with zero bursts in the training or test sets were removed for model fitting.

For the VBN data, CV yielded a somatic subspace of dimension 9, a dendritic subspace of dimension 2 for the base model, and a dendritic subspace of dimension 9 for the hybrid model (Fig. S2a). For the Rodgers data, CV yielded a somatic subspace of dimension 4, a dendritic subspace of dimension 5 for the base model, and a dendritic subspace of dimension 6 for the hybrid model (Fig. S2b).

Evaluating the score difference on both datasets (Fig. 3d,h), we found that the hybrid model beat the null model, demonstrating that dendritic factors beyond those in event rate were required to achieve better predictive performance. Interestingly, for the VBN dataset, the base model performed worse than the null model, whereas for the Rodgers dataset, the two performed equally well. This is consistent with the Rodgers dataset having a larger fraction of burst-coding neurons, as expected given that our analysis was restricted to L5b neurons. In contrast, the VBN dataset included neurons from all cortical layers. All models outperformed the constant burst-probability model (Fig. S3).

For both datasets, we found qualitatively similar results when we analyzed additional sessions, mice, cortical areas, and layers (Fig. S4). However, in some cases the hybrid model did not beat the null model, possibly due to under-sampling of burst coding neurons. Together, these results show that our approach can both identify candidate burst-related structure in real neural recordings and test whether that structure captures the data’s statistics better than event rates alone.

### 5.4 Sensitivity analysis

Given that events were sparse and the mean burst fraction was low for both datasets, we wondered whether our analysis had sufficient power to accurately estimate the somatic and dendritic subspaces. To answer this, we developed a random-matrix model to quantify how subspace error (Eq. 8) depends on the parameters of the base model and the data. See Appendix H for the full theoretical analysis and comparison to MC simulations.

We used our theoretical results (Eqs. S42 and S52), along with the known data and fitted parameters, to estimate the subspace error regime of the model fits in Fig. 3 (see Appendix H.5 for details). We found that somatic and dendritic subspace errors were expected to be reasonably low for both datasets (Fig. S7). Additionally, as expected from theory (Eq. S55), the dendritic subspace error was larger than the somatic subspace error. We also validated our results with MC simulations fit with both the base and hybrid models. Overall, this demonstrates that our analysis in Section 5.3 was not underpowered, and the recovered subspaces are expected to be reasonably accurate.

## 6 Discussion

### Summary

We have introduced MFA, a framework capable of disentangling multiplexed latent factors encoded in events and bursts. Using simulated data, we showed that analyzing firing rates alone muddies these factors, requiring methods that can distinguish between them. Critically, we demonstrated that we can detect the presence of burst codes without requiring detailed knowledge about what is coded in real neural populations.

## Limitations

The application of MFA to neural data is limited by our ability to classify and define bursts. Spike-sorting algorithms can mischaracterize bursts due to their non-standard spike waveform, missing them entirely [63]. MFA is also insensitive to factors encoded in burst length, as it neglects the subsequent spikes in a burst. Burst length is potentially an important variable, as it can encode input slope [64], and short vs. long bursts may have different functional roles [5]. We assumed a burst-detection threshold of 15 ms based on prior work [29, 26], but did not test its impact on our results. Future extensions could refine burst classification, distinguish bursts by length, or infer bursts directly from spike times.

Although a model with dendritic factors predicted bursting fluctuations better than the null model, consistent with some neurons jointly encoding somatic and dendritic factors, other explanations for the hybrid model’s success remain. First, neural compartments may share factors, as multiple pathways target both [65–67]. As a result, bursts may reflect a combination of private and shared factors, potentially explaining why adding a rate component improved predictive performance. Second, a more flexible nonlinear mapping between factors and burst probability may improve performance over the assumed logit-linear mapping [29, 68]. Third, coarse time bins, the omission of factor dynamics, and spike-history effects may lead to some variability being incorrectly attributed to dendritic factors [14, 69]. Future work could explore models with shared and private factors, dynamic factors, and marked point-process spike generation (see Appendix I for model extensions).

Lastly, our sensitivity analysis assumed a homogeneous burst-coding population, along with Gaussiandistributed weights and inference errors. Thus, our theoretical sensitivity estimates may not generalize to more realistic settings (see Appendix H). Future work could relax these assumptions.

## Conclusion

By introducing MFA, we have merged the population and burst coding perspectives together in a principled way, enabling the discovery of burst-related factors. We believe this provides a valuable approach for investigating burst codes, as bursts are thought to encode information relevant to attention, perception, and learning. More generally, our work highlights the importance of considering population structure when investigating temporal codes.

## Data and code availability

All code used to generate synthetic data, fit the models, and analyze the neural data is publicly available on GitHub at https://github.com/ZachFriedenberger/mfa. The neural datasets were previously published and are freely available online [56–58].

## Author contributions

Z.F. contributed to conceptualization, writing, mathematical derivations, simulations, data analysis, and preprocessing of the Rodgers dataset. Y.C. contributed to preprocessing of the VBN dataset. R.N. contributed to conceptualization, writing, and supervision. All authors reviewed, edited, and approved the final manuscript.

### Generative AI usage

Large language models were used to assist with writing and editing code, identifying relevant mathematical identities, and checking intermediate steps in the theoretical derivations underlying the subspace error sensitivity analysis. The authors independently verified all the code and mathematical results.

### Disclosure of funding

This work was supported by an NSERC CGS-D Scholarship, an Ontario Graduate Scholarship, and an NSERC Discovery Grant RGPIN-2024-05162. The authors declare no competing interests.

## A Maximum likelihood estimators for the event and burst emission model

Consider the emission model for events and bursts given by Eq. 2. Given a dataset of event and burst counts, 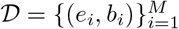, the maximum likelihood estimators for the event rate 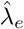 and burst probability 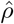 are given by the trial-averaged event-rate and burst fraction,

,
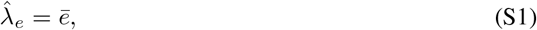

,
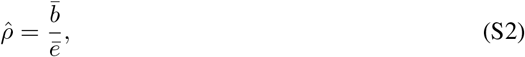

where 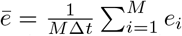 and 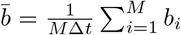 are the trial-averaged event and burst rates. Additionally, the marginal distribution for the number of bursts is given by,

,
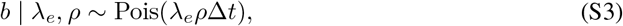

and the estimator for burst rate, *λ*_*b*_ = *λ*_*e*_*ρ*, is given by,

,
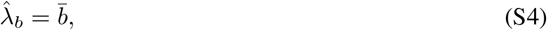

,
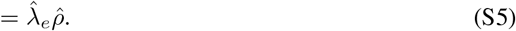

While simple to compute, the application of these estimators is made difficult by two common features of neural data. First, when events are sparse, 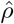 can be undefined. This requires time bins to be large enough such that the trial-averaged event rate is non-zero for each bin, trading off time resolution for numerical stability. Second, these estimators assume that trial-to-trial variability is noise to be averaged away, potentially missing single-trial modulations that are important for behaviour.

## B Distances between linear subspaces

In Figs. 1, S7, and S5, we measured the distance between the true and inferred subspaces using a normalized projection error, which we call the subspace error (Eq. 8). In general, the distance between linear subspaces can be computed using a handful of different metrics, all of which can be expressed in terms of the principal angles between subspaces [70].

The principal angles between two *K*-dimensional subspaces, 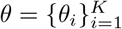, are defined recursively by finding the smallest angle between unit vectors in the two subspaces, then repeating this optimization under the constraint that successive principal vectors within each subspace are mutually orthogonal. Given an orthonormal basis for each subspace, 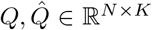, the principal angles can be found using singular value decomposition (SVD),

,
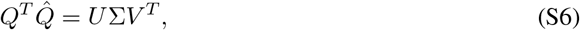

where *U* and *V* are matrices containing the left and right singular vectors, and Σ is a diagonal matrix of singular values such that *σ*_*i*_ = cos *θ*_*i*_.

In terms of principal angles, the subspace error in Eq. 8 becomes,

,
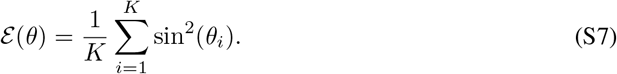

An alternative metric to Eq. S7 is the Grassmann distance *d*_*g*_. The normalized Grassmann distance between subspaces is defined as,

,
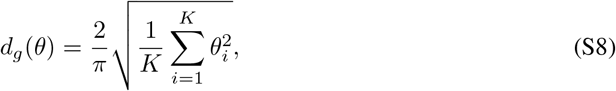

which is also bounded, 0 ≤ *d*_*g*_(*A, B*) ≤ 1, as 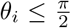. The notion of subspace distance presented here can also be extended to subspaces with different dimensionalities [70].

## C Simulated data

To demonstrate the utility of our MFA framework, we analyzed its performance on both simulated count and spiking data. Here, we explain how the latent factors (or input currents), the factor-loading weights, and the simulated data were generated. All simulated datasets were generated on a ThinkPad P1 Gen 6 laptop with a 13th Gen Intel® Core™ i7-13700H CPU, and required less than 24 hours of execution time.

### Factor loading weights

In all our simulations, we generated random factor-loading weights that defined the somatic and dendritic subspaces. For Figs. 1 and 2, the factor loading weights, 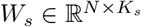 and 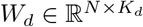, were randomly generated with the Grassmann distance between subspaces held fixed (Eq. S8). For Figs. S5 and S7, the loading weights were simply Gaussian distributed, with 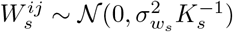 and 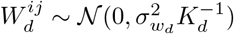.

For factor loading weights with a fixed Grassmann distance, we first generated principal angles by sampling a random direction in the *K*-dimensional angle space and rescaling it to the desired distance. Sampling was repeated until all angles satisfied 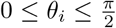. The principal angles were then used to construct random subspaces with a fixed set of principal angles.

We first sampled an orthonormal somatic loading basis, 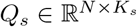, and an orthogonal complement 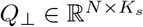. Assuming *K*_*d*_ *K*_*s*_, the dendritic loading basis was then constructed by rotating the somatic basis into the orthogonal complement,

,
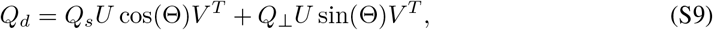

where cos(Θ) and sin(Θ) are *K*_*d*_ × *K*_*d*_ diagonal matrices of the cosines and sines of the principal angles. 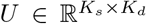 and 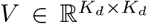 are random orthonormal matrices that determine the directions in the somatic subspace that are rotated and the basis of the dendritic subspace, respectively. These matrices were generated by performing SVD on a random Gaussian matrix, 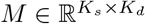. For one-dimensional subspaces, *K*_*s*_ = *K*_*d*_ = 1, Eq. S9 reduces to,

,
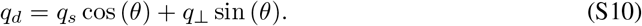

Lastly, we scaled the orthonormal bases to set the variance of the factor loadings, 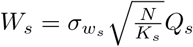 and 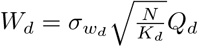. Where the factor of 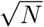 was added so that the scale of the loading weights would be similar to the case when there was no fixed Grassmann distance (unnormalized), and the factor of 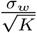 sets the variance.

### One-dimensional latent factor data

For Fig. 1, one-dimensional somatic and dendritic latent factors were generated from a zero-mean Gaussian process,

,
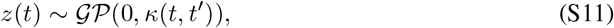

where *κ* is a squared exponential kernel with unit variance,

,
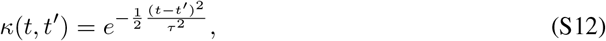

and temporal smoothness governed by a timescale *τ*. The total length of the latent factors was chosen to be 500 time steps of size Δ*t* = 1 (i.e., *M* = 500 samples). The timescale was chosen to be *τ* = 10 for both the soma and the dendrite.

The factor loadings for the one-dimensional subspaces were randomly generated with a fixed angle between them, as described above (Eq. S10). The parameters for the loadings were, *N* = 200, and 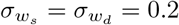. The offsets defining the baseline event rate and burst probability were identical across all neurons and set to *µ*_*s*_ = 2 and *µ*_*d*_ = 0.

Observed counts were generated using Eqs. 2 and 3, treating the time axis of the latent factors as independent samples. The total number of spikes was then calculated using *a* = *e* + (*n* − 1)*b*, where the number of spikes per burst was set to *n* = 3. This procedure resulted in the data matrices, *E, B, A* ∈ ℝ^*N* ×*M*^. In Fig. 1b, a single simulated dataset was used as an example, while in Fig. 1c-d, 100 simulated datasets were used for each burst power and angle pair.

### Spiking population data

The simulated spiking datasets were generated from a population of two-compartment model neurons, which were modelled after thick-tufted pyramidal neurons. See Naud & Sprekeler for the full model [29]. We used the same neuron parameters as in their work and report only parameter changes and differences in the input currents.

We generated data for multiple datasets, varying the number of neurons with the dendrite intact and silenced (Fig. 2). For neurons with an intact dendrite, the coupling between compartments was set to *g*_*s*_ = 1300 pA. When the dendrite was silenced, the coupling was set to zero, *g*_*s*_ = 0 pA, transforming the two-compartment model into an adaptive leaky-integrate-and-fire model. Additionally, we increased the amount of dendritic adaptation, 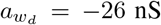, to reduce the mean burst length. In all simulations, populations contained *N* = 50 neurons. Each neuron’s somatic and dendritic compartment received an input current composed of a shared, private, and noise component,

,
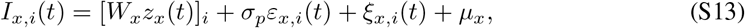

where *x* ∈ {*s, d}* and *i* is the neuron index. The shared latent factors, *z*_*x*_(*t*), were constant over an interval of 250 ms with an amplitude sampled from a standard normal distribution. The somatic and dendritic factor loadings, *W*_*s*_ and *W*_*d*_, defined 3-dimensional subspaces, *K*_*s*_ = *K*_*d*_ = 3. Loadings were sampled with a fixed distance between them, *d*_*g*_ = 0.5, as described previously. The standard deviation of the loadings was set to 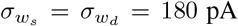. Each compartment and neuron also received a private time-varying input, *ε*_*x,i*_(*t*), that was constant over an interval of 250 ms with an amplitude sampled from a standard normal distribution, scaled by *σ*_*p*_ = 120 pA. The background noise currents, *ξ*_*x*_(*t*), were sampled from an Ornstein-Uhlenbeck (OU) process for each neuron. The parameters of the noise current were set to, 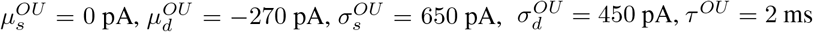. Lastly, the baseline currents were set to *µ*_*s*_ = *µ*_*d*_ = 250 pA for each neuron.

Neural dynamics were integrated using an exponential integration scheme with an integration time step of 0.1 ms. Each simulation produced data for both a training and a test set. The training set consisted of 500 trials, each lasting 5250 ms. The first 250 ms of each trial were discarded to remove transient effects at the beginning of the simulation. Spike times were recorded upon crossing a threshold and then classified into events and bursts. Counts were obtained over the 250 ms windows in which the latent factors were constant. This resulted in a total of *M* = 10, 000 samples for the training set. Test datasets were generated with 100 trials, yielding a total of *M* = 2, 000 test samples.

### Synthetic experimental data

The simulated experimental data that was fit in Fig. S7 was generated directly from the base and hybrid MFA models (Eqs. 2, 3, and 5). The number of neurons *N* and samples *M* were identical to the real data. The offset parameters for each neuron, *µ*_*s*_ and *µ*_*d*_, and the number of latent factors, *K*_*s*_ and *K*_*d*_, used for data generation were identical to what was inferred from the real data. As stated above, factor loadings were Gaussian distributed, with 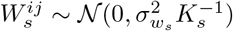 and 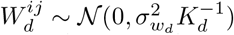. The variances of the factor loadings, 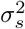 and 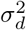, were estimated from the inferred factor loadings, as described in Section 5.4. For the hybrid model simulations, the neuronspecific dendritic weights *w*_*d*_ were identical to those inferred from the real data. This produced datasets with the same number of samples and a similar population structure to the real data.

## D Experimental data

The data in Fig. 3a-d, Fig. S2a, Fig. S3a, and Fig. S4a-c are from the Allen Institute’s openly available Visual Behavior Neuropixels (VBN) dataset [56, 57]. The data in Fig. 3e-h, Fig. S2b, Fig. S3b, and Fig. S4d-f are from the Rodgers dataset [58]. For a full description of the experimental setup and extracellular recordings, please see the corresponding white paper for the VBN dataset and the data paper for the Rodgers dataset. Here, we provide additional details on each behavioural task, data preprocessing, and the subset of each dataset analyzed.

### D.1 Visual Behavior Neuropixels dataset

#### Data source and license

We used version v0.5.0 of the VBN dataset. The dataset is archived on DANDI (Dandiset 000713) under CC BY 4.0.

#### Behavioural task

Mice performed a visual change-detection task in which they were presented with a sequence of natural images (250 ms), interleaved with a grey screen (500 ms). The same image was repeated until it was randomly replaced with another image (change image), after which the mice were required to report the change by licking a spout for a water reward within 750 ms. Trials in which the mouse received the reward are called “hit” trials (Fig. 3b).

Mice performed the task in two sessions: a familiar session comprising 8 images seen during training, and a novel session comprising 6 new images and 2 familiar images. Prior analysis of this dataset has shown that mice perform worse on the familiar images in the novel session, and that this is accompanied by an increase in the dimensionality of the population activity as measured using firing rates [71].

#### Data preprocessing

Well-isolated units were selected using the spike-sorting metrics provided with the dataset. A detailed summary of each of these metrics is provided by Siegle et al. [72]. We chose a subset of these metrics and set their thresholds to minimize the number of contaminated units. First, we filtered out all units whose templates were classified as noise, and then we used the default AllenSDK metrics but decreased the tolerance for ISI violations. Specifically, we kept only units with an overall firing rate > 0.1 Hz, an Amplitude cutoff < 0.1, a Presence ratio > 0.9, and ISI violations = 0. This drastically reduced the dataset size, resulting in approximately 20-80 visual cortical neurons per session.

#### Data analyzed

Mice with the ID numbers 599294, 574082, and 530862 were chosen for analysis because they had more than 30 neurons per session and a low event sparsity. The analysis presented in Section 5.3 corresponds to the novel session for mouse 599294. The results for the familiar session and the other two mice are shown in Fig. S4a-c. Mice often had high task accuracy in the familiar session, which dropped during the novel session. Mouse 599294 had an accuracy of 86% on the familiar session and 63% on the novel session.

### D.2 Rodgers dataset

#### Data source and license

We used the Rodgers dataset from DANDI (Dandiset 000231, draft version last updated in June 2023), under CC0 1.0.

#### Behavioural task

Mice performed a whisker-based shape-discrimination task, distinguishing concave from convex objects in darkness. At the beginning of each trial, a linear servo motor moved either shape towards the whiskers and stopped at one of three distances (close, medium, or far). The shapes stopped 1.2-1.6 s after they began moving. Importantly, mice needed to actively whisk to sense the object, even in the close condition. At 2 s, a response window opened, where the mouse could lick left to report a concave shape or right to report a convex shape. The first lick in the response window was recorded as the choice. Correct choices were rewarded with water, while an incorrect choice led to a 9 s timeout.

#### Data preprocessing

Spike sorting was originally conducted using a combination of Kilosort and manual curation, and similar metrics to those above were used for quality control. We used the anatomical location (depth within cortex) provided with the dataset to select units that were labelled as L5b or deep neurons (L5a, L5b, and L6). Additionally, we restricted our analysis to putative excitatory neurons. Non-random trials and those with optogenetic stimulation were dropped from the analysis. We analyzed ±3 s around the choice time and verified that no trials overlapped.

#### Data analyzed

Mice with the subject IDs KM131, KF132, and CR221 were used in our analysis. For each mouse, we analyzed two sessions corresponding to different recording days (Fig. S4). The data shown in Fig. 3e-h correspond to session 20180118 from mouse KM131. The results for an additional session and the other two mice are shown in Fig. S4d-f. In general, mice performed the task well. For example, mouse KM131 had an accuracy of 87% on session 20180118 and 78% on session 20180119.

## E Model inference details

The simulated and experimental datasets were fit using the inference procedures introduced in Section 4.1. Here, we provide additional implementation details and specify which procedure was used for each analysis reported in Section 5. All model fitting was performed on a ThinkPad P1 Gen 6 laptop with a 13th Gen Intel® Core™ i7-13700H CPU, and required less than 24 hours of execution time.

### One-dimensional latent factor data

For Fig. 1, the Gaussian-approximate EM procedure described in Section 4.1 was used for inference. This was implemented using the FactorAnalysis class from scikit-learn [73]. In Fig. 1b-c, the number of latent dimensions was fixed and not inferred from the data. The FA model was fit using the total number of spikes and 2 factors, whereas the MFA base model was fit using events and bursts, with 1 somatic and 1 dendritic factor. For Fig. 1d, the dimensionality was chosen using the K-fold CV approach described in Section 4.2, with 5 folds. For this simulation, we evaluated models with up to 5 dimensions and selected the one with the largest mean score over folds.

### Spiking population and experimental data

For Figs. 2, 3, and S7, the VI procedure described in Section 4.1 was used for inference. In all cases, we optimized the objective with Adam for *T* = 5000 iterations, computing gradients over the full training set at each iteration [74]. The learning rate decayed exponentially, 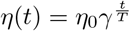, where *η*_0_ = 10^−2^ and *γ* = 10^−4^.

Model dimensionality was again selected using 5-fold cross-validation (CV). To reduce the influence of inter-fold variability, we computed the difference in CV score between each candidate model and the simplest model in its class within each fold, then averaged these differences across folds. We refer to this quantity as the CV score gain, as it measures the improvement in predictive performance relative to a simple baseline model. For the event and burst components of the base model, the baseline models had *K*_*s*_ = 1 and *K*_*d*_ = 1, respectively. For the burst component of the hybrid model, the baseline had *K*_*d*_ = 0, corresponding to the null model. To select dimensionality, we first identified the number of dimensions that maximized the mean CV score gain, then selected the smallest number of dimensions whose mean gain was within one standard error of this maximum. For each model, we first estimated the dimensionality of the somatic subspace, then held it fixed while estimating the dendritic dimensionality. We found that this procedure most accurately identified the knee of the score-versus-dimension curve in simulated spiking data.

For the simulated spiking datasets, we evaluated models with up to 6 somatic and 9 dendritic dimensions. Figure S1 shows the results of this procedure, with the selected dimensions indicated by black stars. In all cases, the true somatic dimensionality was correctly recovered. When all dendrites were intact, both models also recovered the correct dendritic dimensionality (Fig. S1a). However, when all dendrites were silenced, the hybrid model correctly selected *K*_*d*_ = 0, whereas the base model selected *K*_*d*_ = 3 (Fig. S1b). These spurious dimensions arose because the base model used additional dendritic factors to approximate the rate-dependent contribution to bursting. Similarly, when the population contained a mixture of intact and silenced dendrites, the hybrid model recovered the correct dendritic dimensionality, whereas the base model overestimated it by using additional factors to capture both rate-dependent bursting and independent dendritic input (Fig. S1c).

For the experimental datasets, we evaluated models with up to 15 somatic and 15 dendritic dimensions, or up to the number of recorded neurons when fewer than 15 were available. The dimensionality estimation results corresponding to the data in Fig. 3 are shown in Fig. S2, with the VBN and Rodgers datasets shown in Fig. S2a and Fig. S2b, respectively. CV was not performed for the synthetic experimental data, as the dimensionality was fixed to that inferred from the corresponding experimental dataset.

## F Independent model comparison

For each dataset, we additionally compared the base, hybrid, and null models to an independent model in which each neuron had a burst probability that was constant across samples (Section 3.2). We found that all the models performed better than the independent model for both the VBN (Fig. S3a) and Rodgers datasets (Fig. S3b). This demonstrates that the bursts are explained better when taking population structure into account.

## G Additional recording sessions

As mentioned in Sections 5.3 and 4.3, we analyzed additional recording sessions for each dataset. Figure S4 shows the score difference for the base-null and hybrid-null model comparisons for five additional sessions per dataset. Figure S4a shows the familiar session from the same mouse shown in Fig. 3, while Figs. S4b-c show new mice from the VBN dataset. Figure S4d shows another session of the Rodgers dataset for the mouse in Fig. 3, while Figs. S4e-f show new mice, with data from deep neurons (L5 and L6) recorded from barrel C3.

**Figure S1:**
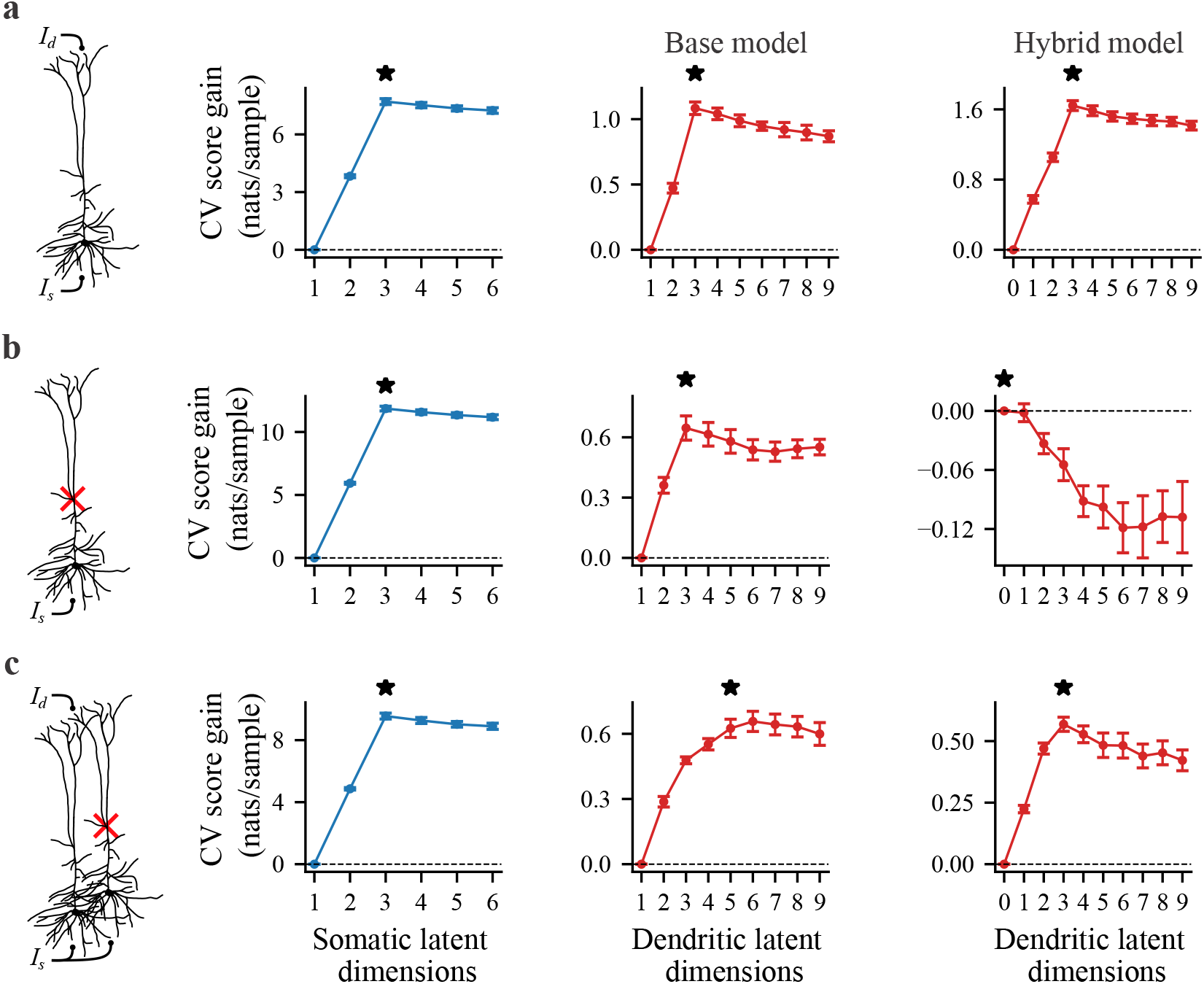
Subspace dimensionality estimation for the simulated spiking data presented in Fig. 2. **a**, Intact, **b**, silenced, and **c**, mixed intact and silenced dendrite simulations. These correspond to populations with fractions of (1.0, 0.0, and 0.5) intact dendrites, respectively. The cross-validated (CV) score gain averaged over folds is plotted as a function of the number of somatic and dendritic latent dimensions. Somatic dimensionality estimation (left) and the dendritic dimensionality estimation for both the base (middle) and hybrid (right) models. Black stars denote the selected dimensionality. Error bars represent ±1 s.e.

## H Subspace error theory and sensitivity analysis

To determine how well the somatic and dendritic subspaces could be recovered in practice, we used theory to quantify how the expected subspace error given by Eq. 8 scales with the model and data parameters. We developed a random-matrix-model formalism that allowed us to obtain the error scaling in the low-error regime. Assuming small Gaussian errors on the inferred loading weights, we showed that the expected subspace error can be computed analytically given the variance of the Gaussian errors (H.1). We estimate these Gaussian errors using the complete-data Fisher Information Matrix (FIM) in Section H.2, resulting in a lower bound on the expected somatic and dendritic subspace error (Section H.3). While the real errors on the inferred parameters are most likely not independent and identically distributed, nor Gaussian, we verified that our theory matches the error obtained using MC simulations well over a large range of the parameters (Section H.4). Lastly, as described in Section 5.4, we used our theory and simulations to estimate the subspace error of the experimental data fits (Section H.5).

**Figure S2:**
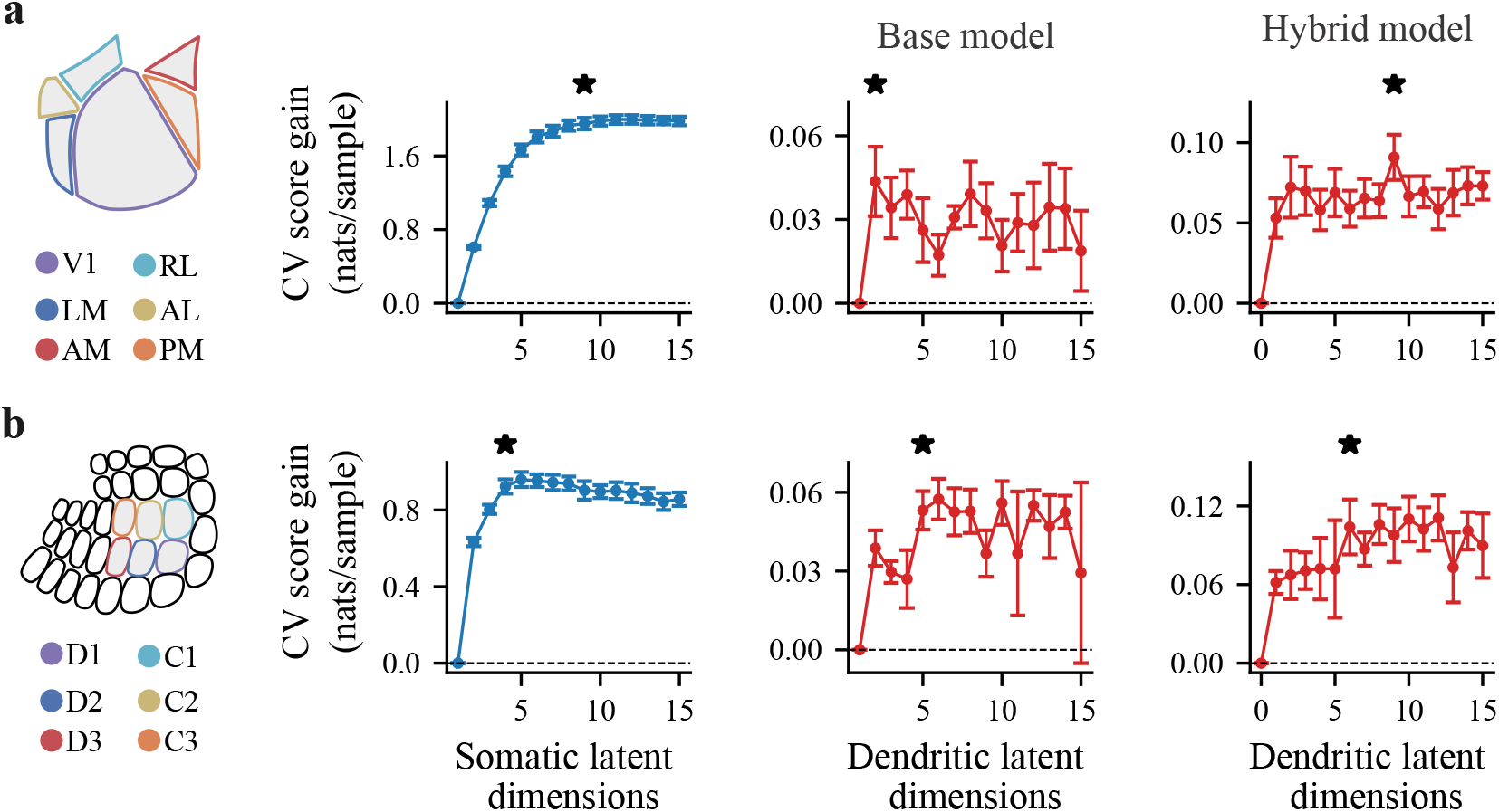
Subspace dimensionality estimation for the datasets and sessions presented in Fig. 3. **a**, VBN dataset. **b**, Rodgers dataset. The cross-validated (CV) score gain averaged over folds is plotted as a function of the number of somatic and dendritic latent dimensions. Somatic dimensionality estimation (left) and the dendritic dimensionality estimation for both the base (middle) and hybrid (right) models. Black stars denote the selected dimensionality. Error bars represent ±1 s.e.

**Figure S3:**
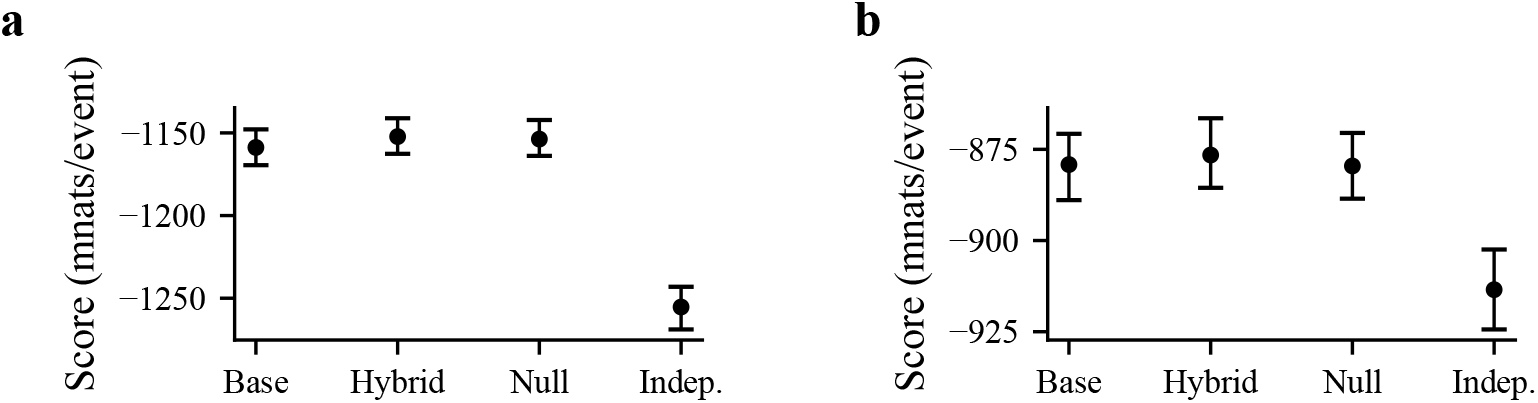
Comparison of models trained and evaluated using the *in vivo* neural datasets shown in Fig. 3. **a**, VBN dataset. **b**, Rodgers dataset. The mean predictive log-likelihood per event of the base, hybrid, null, and constant burst-probability (Indep.) models are compared. Error bars are 95% confidence intervals obtained via bootstrapping over test samples.

### H.1 A random matrix model for subspace error

We considered a random matrix model for the estimated subspace. As only perturbations orthogonal to the true subspace increase the error, we assumed that the estimated subspace 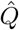 is given by,

,
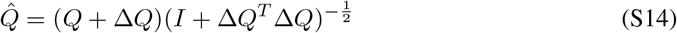

where *Q* is an orthonormal matrix of the true subspace, *Q*^*T*^ *Q* = *I*_*K*_, Δ*Q* is a small random perturbation orthogonal to the true subspace, *Q*^*T*^ Δ*Q* = 0, and the term 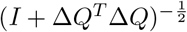 ensures that 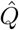 is orthonormal, 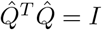.

Using this parametrization of the estimated subspace, we found the subspace error to first order in Δ*Q*. First, we expanded Eq. S14 to first order,

,
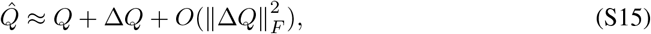

and then plugged Eq. S15 into Eq. 8. After discarding higher-order terms and using the orthonormal and orthogonality conditions, we found that the subspace error reduces to,

**Figure S4:**
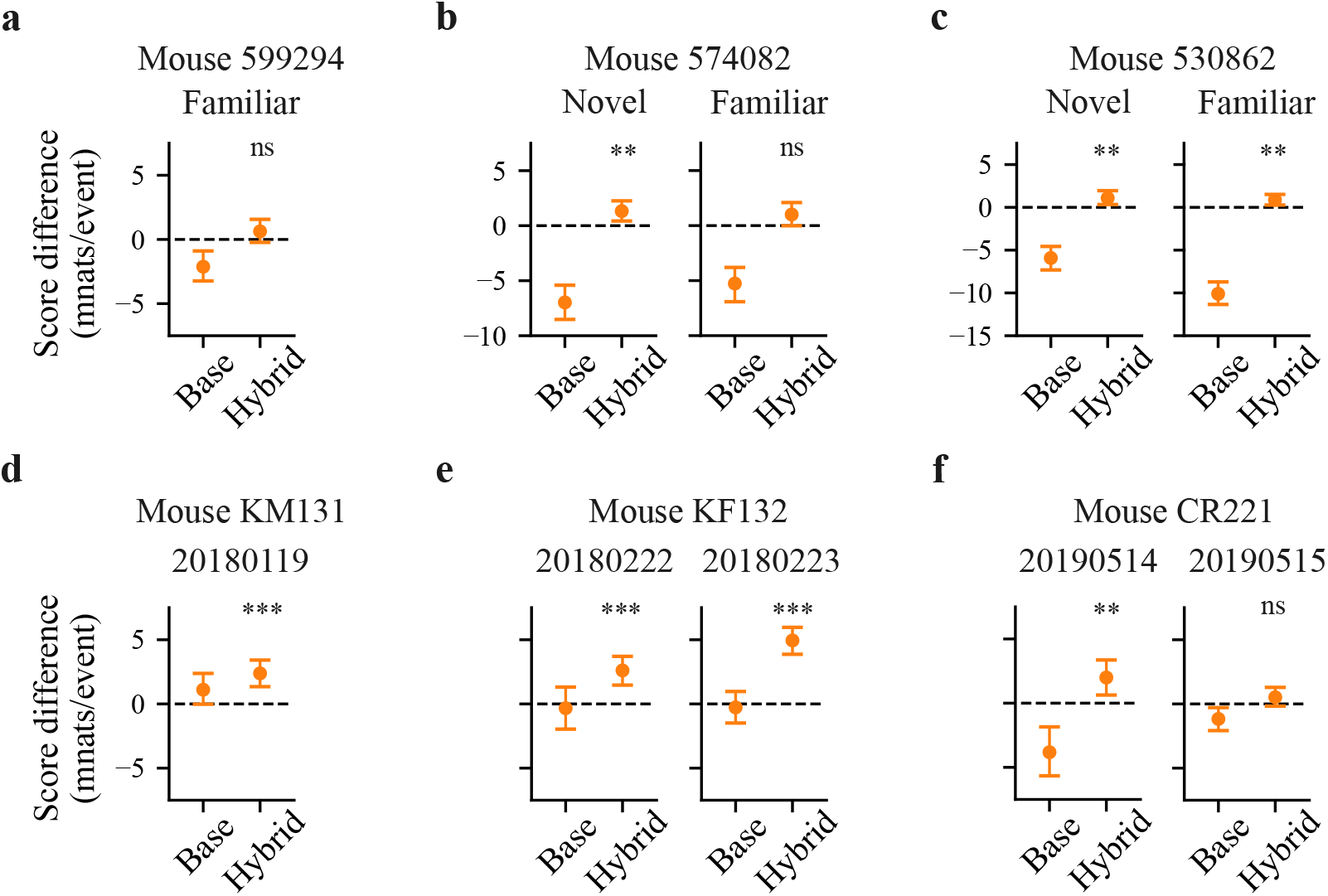
Analysis of additional recording sessions. The difference in mean predictive log-likelihood per event (Score difference) for base-null and hybrid-null. **a-c**, VBN dataset. **a**, The familiar session for the mouse shown in Fig. 3d. **b-c**, Novel and familiar sessions from additional mice. **d-f**, Rodgers dataset. **d**, An additional recording session for the same mouse shown in Fig. 3h. Similarly, the recording was in barrel D2, and only L5b neurons were analyzed. **e-f**, Additional mice and recording sessions from the Rodgers dataset. The recordings were in barrel C3, and only deep neurons (L5a, L5b, and L6) were analyzed. Error bars show the bootstrapped 95% confidence interval; ** and *** denote *p* ≤ 0.01 and *p* ≤ 0.001 for a two-sided bootstrap test of a score difference of zero; ns denotes not significant.

**Figure S5:**
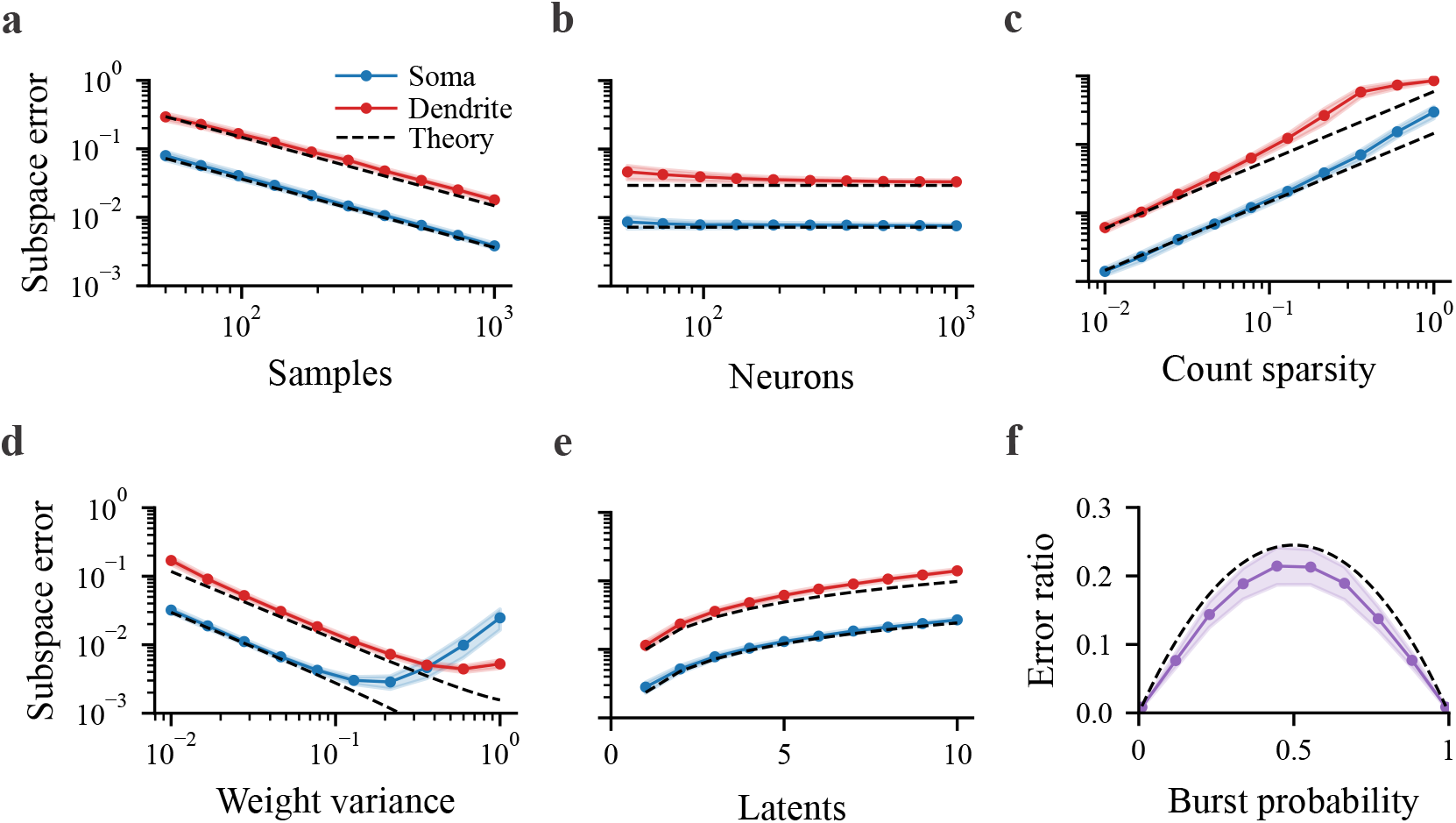
Sensitivity analysis of latent subspace recovery using MFA. **a-e**, Simulated somatic (blue) and dendritic (red) subspace error as a function of model and data parameters. **f**, Ratio of the somatic to dendritic subspace error (purple) as a function of baseline burst probability. Coloured lines are the mean subspace error over repeated simulations, with the shaded region showing one standard deviation. Black dashed lines show the theoretically expected error. For all our simulations, one parameter is varied while the others are fixed, except for **d**, where the somatic error is shown as a function of the somatic weight variance for fixed dendritic weight variance, and vice versa for the dendritic error.

**Figure S6:**
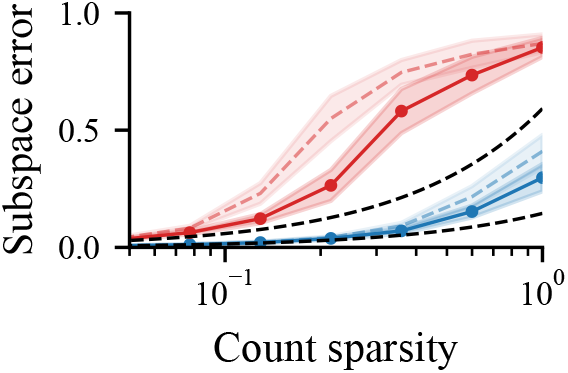
Subspace error obtained by the different inference procedures as a function of event count sparsity 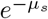. Simulated somatic (blue) and dendritic (red) subspace error. Gaussian-approximate EM (dashed coloured lines) and VI inference (solid coloured lines). Black dashed lines show the theoretically expected error. VI recovers the dendritic subspace better than Gaussian-approximate EM when event counts are sparse. The VI and theory curves are identical to Fig. S5c, but the axes have been modified.

**Figure S7:**
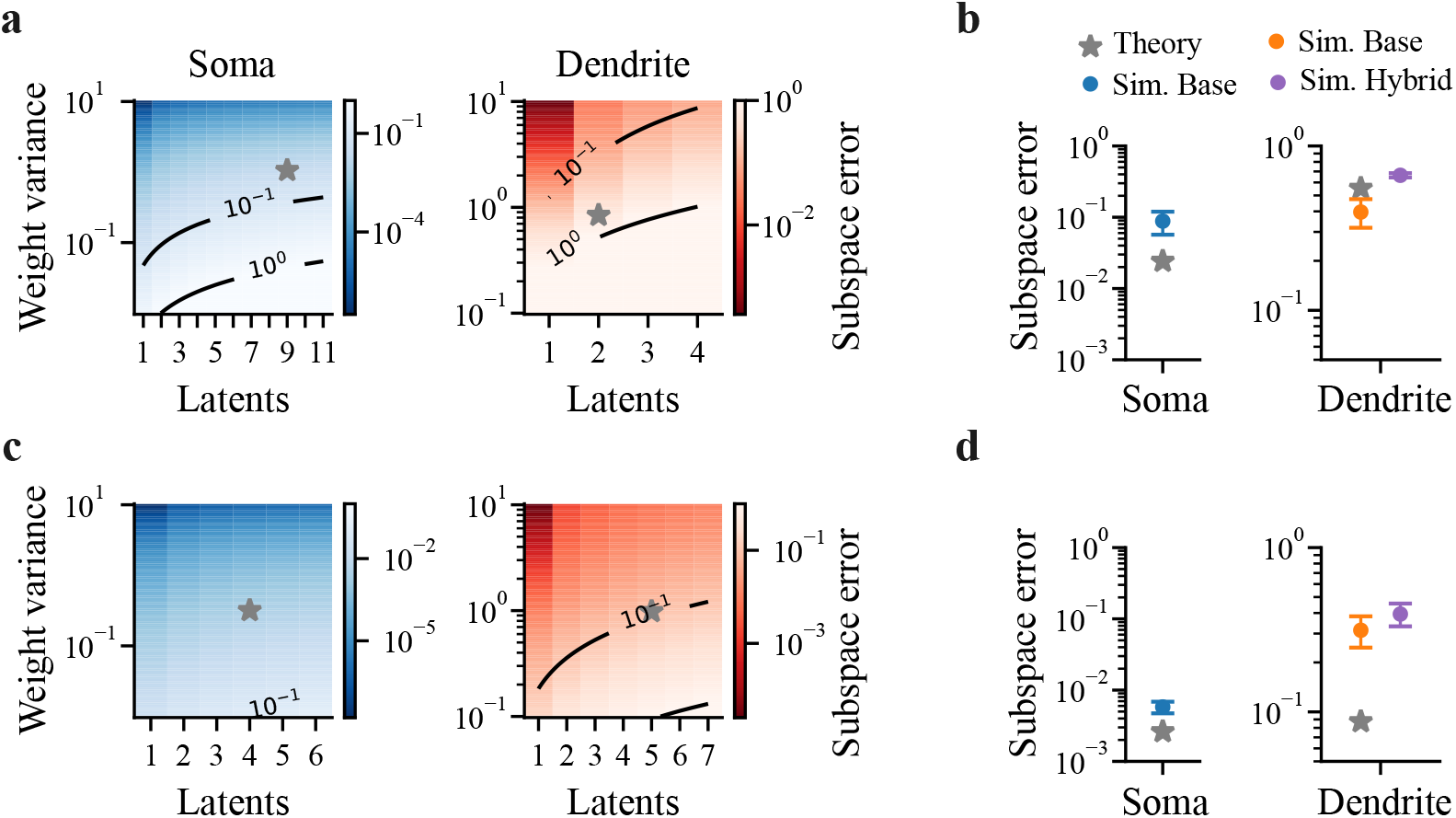
Sensitivity analysis. The data and model fits are identical to those shown in Fig. 3. **a-b**, VBN dataset. **a**, Heatmaps of the expected somatic and dendritic subspace error for the base model as a function of the weight variance and number of latent dimensions, obtained using theory. Black lines show the level sets for different errors. Point estimate of the weight variance and number of dimensions (grey star). **b**, Expected somatic (left) and dendritic (right) subspace error. The theoretical value of the error at the point estimate (grey star), the simulated error for the base event model (blue), the base model (orange), and the hybrid model (purple) are shown. **c-d**, Same as **a-b** but for the Rodgers dataset. Error bars are ± 1 s.e. in the mean across 5 repeated simulations with randomly generated weights.

,
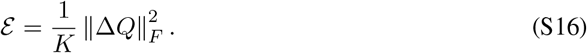

This can be rewritten using the trace definition for the Frobenius norm,

,
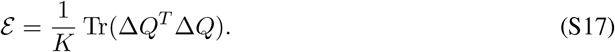

Note that the version of 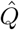 used in the expansion given by Eq. S15 is non-orthonormal, and we will want to find a similar 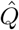 to use Eq. S17.

We then related the random errors in the estimated factor loading weights Δ*W* to the random orthogonal perturbation Δ*Q*, by assuming that the estimated loadings are given by,

,
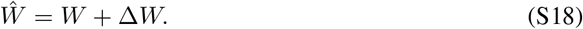

Where *W* are the true loadings and Δ*W* is a small Gaussian perturbation, such that each element has a random error, Δ*W*_*ij*_ ~ *N* (0, *σ*^2^). The variance of this perturbation, *σ*^2^, therefore, characterizes the error in estimating each element of the factor loading weight matrix.

Using a QR decomposition of the true loadings, *W* = *QR*, we obtained an orthonormal matrix *Q* for the true subspace and an upper triangular matrix *R*. Using this decomposition, we expressed the estimated loading weights in a useful orthonormal basis,

,
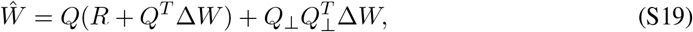

where *Q*_⊥_ ∈ ℝ^*N* ×(*N* −*K*)^ is the orthogonal complement of *Q*. Here we have used the fact that 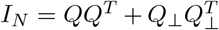. By right multiplying both sides of Eq. S19 by (*R* + *Q*^*T*^ Δ*W*)^−1^ we obtained,

,
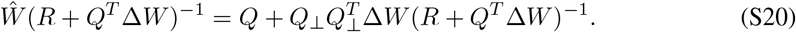

Importantly, this operation does not change the column space of the left-hand side, which is identical to the column space of the estimated subspace 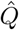 and loadings *Ŵ*. Therefore, by expanding the right-hand side and dropping higher-order terms, we found,

,
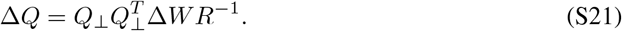

Using Eq. S21 as the orthogonal perturbation to the true subspace and plugging it into Eq. S17 gave,

,
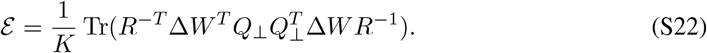

Taking the expectation over the random Gaussian perturbation Δ*W*,

,
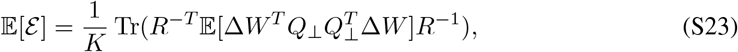

,
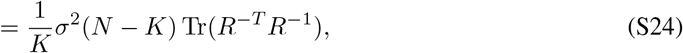

where we have used the fact that the trace and expectation operators are exchangeable. Lastly, QR decomposition of the true subspace again results in,

,
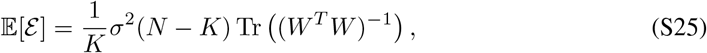

which depends on the variance of the errors *σ*^2^, the size of *W*, as well as its singular values 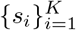, as 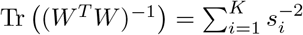.

While straightforward to compute, Eq. S25 requires that properties of the true factor loading weights be known exactly. Therefore, to proceed further, we assumed that the true weights themselves are random matrices, 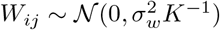. Taking the expectation over the true loading weights and computing the trace gives,

,
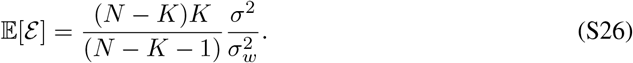

Where we have used the fact that E Tr (*W*^*T*^ *W*)^−1^ can be computed by exchanging the order of the trace and expectation, and recognizing that (*W*^*T*^ *W*)^−1^ is distributed according to the inverse Wishart distribution with *N* degrees of freedom, 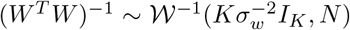. If *N* is large compared to *K*, Eq. S26 simplifies to,

,
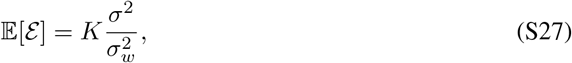

which can be interpreted as an inverse signal-to-noise ratio. Note that this is only valid when 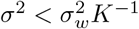, as the subspace error is bounded ℰ ≤ 1. Additionally, the first-order approximation requires that *σ*^2^ be small. Using MC simulations, not reported here, we find that Eq. S27 is most accurate when 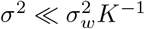.

### H.2 Estimating the error on the inferred parameters

To estimate the error on the inferred parameters *σ*^2^ used in Eq. S27, we used the mean variance of the inferred loadings,

,
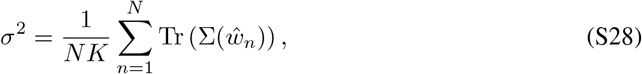

where Σ(*ŵ* _*n*_) is the covariance matrix for the inferred neuron-specific factor loadings, 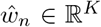. In general, the covariance matrix in Eq. S28 can be approximated using the inverse of the FIM, 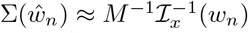, where *M* is the number of samples, ℐ_*x*_ is the FIM for a single sample, and *w*_*n*_ is the true neuron-specific loadings [75]. For ease of notation, we drop the neuron subscript *n*.

For a model with marginal log-likelihood, log *p*(*x* | *w*), if the second derivative with respect to the loadings exists, the observed FIM is given by,

,
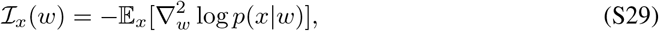

where *x* is the observed data and *w* are the loadings for a single neuron. Unfortunately for MFA, the marginal log-likelihood is intractable and therefore Eq. S29 is also intractable. While the FIM can be computed for latent variable models using numerical methods, we instead use the complete-data log-likelihood as a simple analytical approximation.

For a latent variable model with complete-data log-likelihood, log *p*(*x, z* | *w*), the observed FIM is given by the chain rule,

,
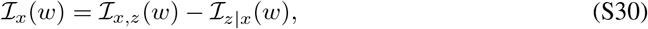

where ℐ _*x,z*_(*w*) is the complete-data term and ℐ_*z*|*x*_ (*w*) is the missing information term. The completedata term is the Fisher information when both *x* and *z* are observed and is given by,

,
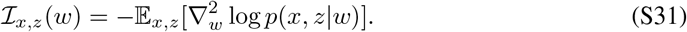

While the missing information term is the amount of information lost due to *z* being a latent variable,

,
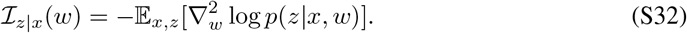

We used the inverse of the complete-data Fisher information as an optimistic approximation for the covariance of the loadings, 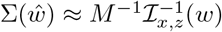. Therefore, the error on the inferred loadings was taken to be,

,
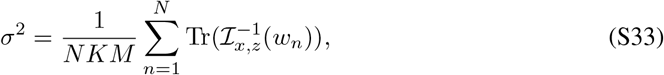

where 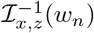 is the inverse of the complete-data FIM evaluated at the true neuron-specific loadings *w*_*n*_ ∈ ℝ^*K*^. As the complete-data FIM ℐ_*x,z*_(*w*) is greater than or equal to the observed Fisher information ℐ_*x*_(*w*), we expect to underestimate the true parameter variance and thus the subspace error. This is verified in Fig. S5a-e, where our theory appears as a tight lower bound on the simulated subspace error.

### H.3 Derivation of the expected somatic and dendritic subspace error

For our MFA model, the factor loadings only enter in the conditional likelihood, *p*(*x, z*|*w*) = *p*(*x*|*z, w*)*p*(*z*). Therefore, the complete-data FIM from Eq. S31 simplifies to,

,
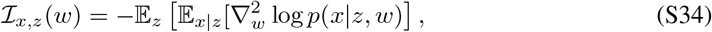

where the conditional expectation is taken with *z* fixed. As the log-likelihood of our model decomposes into,

,
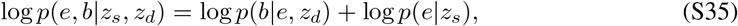

the complete-data FIM for the somatic and dendritic subspace is given by,

,
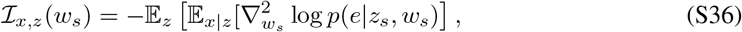

,
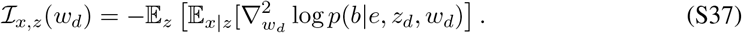

In the following, we derive the complete-data FIM and the expected error for each subspace. We assume that the true loadings are normally distributed, such that 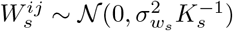 and 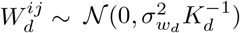.

#### Somatic subspace error

We find that calculating the second derivative with respect to *w*_*s*_ in Eq. S36 gives,

,
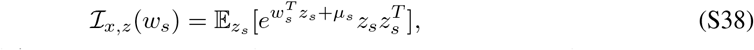

where we have absorbed Δ*t* into *µ*_*s*_. Evaluating the Gaussian expectation we find,

,
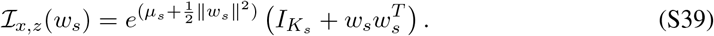

Taking the inverse then gives,

,
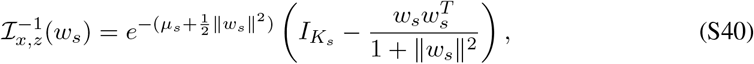

where we have used the Sherman-Morrison inverse for the 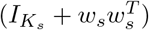 term. The trace is then given by,

,
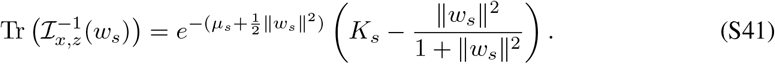

Using the fact that we assumed the true loadings are Gaussian distributed with fixed variance, we have 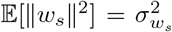. Combining Eq. S41 and S33, and reintroducing the neuron index *n*, the expected subspace error is given by,

,
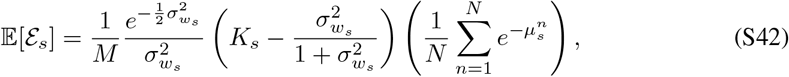

where we have approximated the nonlinear function of the squared loading norms by evaluating it at their mean. This equation can be understood as a product of three terms: a loading-dependent term, the mean event sparsity, and the inverse number of samples.

#### Dendritic subspace error

We find that calculating the second derivative with respect to *w*_*d*_ in Eq. S37 gives,

,
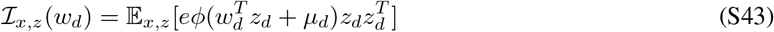

,
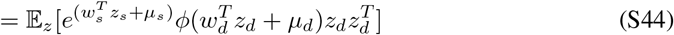

,
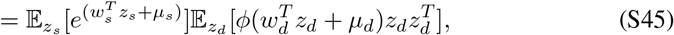

where *ϕ*(*x*) = sig(*x*)(1 − sig(*x*)), sig(*x*) = (1 + *e*^−*x*^)^−1^, and we have absorbed Δ*t* into *µ*_*s*_. We find that the first expectation with respect to the somatic latent factors is,

,
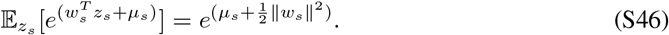

The second expectation with respect to the dendritic latent factors, however, needs to be approximated. Here, we use Stein’s lemma to simplify the expectation [76]. For a scalar function *f* (*z*) of the latent factors, applying Stein’s lemma twice gives the identity,

,
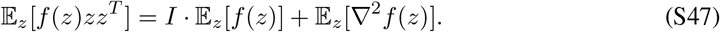

*d*

Second, we let 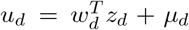 and recognize that the distribution of *u*_*d*_ is given by, *u*_*d*_ ~ *N* (*µ*_*d*_, ∥*w*_*d*_∥ ^2^). Rewriting the expectations in Eq. S47 in terms of the function *ϕ*, the latent factors *z*_*d*_, and the one-dimensional variable *u*_*d*_ gives,

,
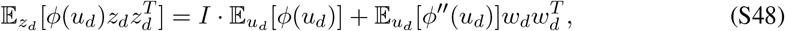

where *ϕ*^′′^ is the second derivative of the scalar function. We then approximated the one-dimensional Gaussian expectations using the delta rule to first order, E[*f* (*u*)] ≈ *f* (E[*u*]). Approximating the expectations to second order did not make much of a difference to our final results.

Putting everything together, the complete-data FIM is given by,

,
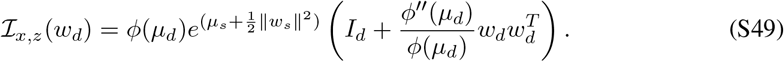

Letting *γ*(*x*) = *ϕ*^′′^(*x*)*/ϕ*(*x*) = 1 − 6sig(*x*) + 6sig^2^(*x*) and taking the Sherman-Morrison inverse gives,

,
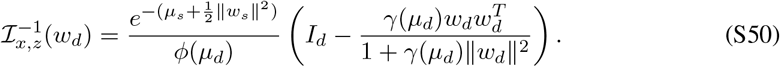

Taking the trace, we find,

,
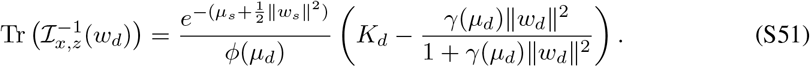

Similar to the somatic subspace error, we use the fact that 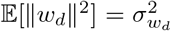. Combining Eq. S46, S51 and S33, and reintroducing the neuron index *n*, the expected subspace error is given by,

,
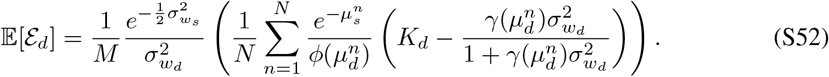

Compared to the expected somatic subspace error, the dendritic subspace error is not easily interpretable in this form. Therefore, we consider the case when the baseline offset parameters are constant across neurons.

#### Simplified subspace errors

Equations S42 and S52 can be simplified by assuming the *µ*_*s*_ and *µ*_*d*_ are constant across neurons. In that case, the expected somatic subspace error can be written as,

,
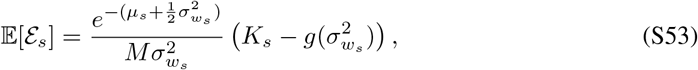

where 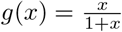. Similarly, the expected dendritic subspace error can be written as,

,
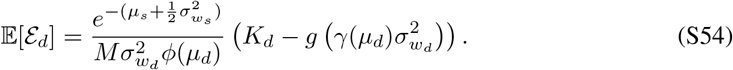

We analyze these equations numerically and compare with MC simulations in the next section.

### H.4 Comparison with Monte Carlo simulations

To verify our theoretical results in Eqs. S53 and S54, we compared the theoretically predicted error with the error obtained using MC simulations. For each MC sample, the true loading weights were sampled as above, and counts were generated from the MFA model. Each simulated dataset was fit using either the Gaussian-approximate EM or VI inference procedure, depending on the event count sparsity. Estimated loading weights *Ŵ* were compared to the true loading weights *W* using the subspace error (Eq. 8). We found that Eqs. S53 and S54 characterized the subspace errors well over a wide range of parameters (Fig. S5), accurately predicting the mean simulated error in the small error regime (30 MC sampled datasets per point).

While we made assumptions about the structure of the true loadings, this analysis gives us a better understanding of how these different subspaces can be estimated in practice. In terms of parameters that can be experimentally controlled, Fig. S5a-b shows that the subspace error decreases with the number of samples *M* and is constant in the number of neurons *N*. Figure S5c shows that subspace error increases with the event count sparsity 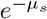. We found that when fitting data with high event count sparsity, the VI inference procedure performed much better than the Gaussian-approximate method (Fig. S6). Taken together, this implies that in sparse spiking data, there is a trade-off between the size of the bin used for counting events and the number of samples.

In terms of the true loadings, Fig. S5d-e shows the subspace error as a function of the loading variance parameter 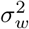 and number of latent dimensions *K* for each subspace. Error increases with the number of latent dimensions; however, this is highly dependent on the scaling we used in our simulation for the true loadings. When the weights are not scaled by the number of dimensions, it is expected to scale differently. For small weight variances, the error decreases and matches the theory closely, before increasing and deviating from the theory at larger values.

Interestingly, we found that the somatic subspace error was smaller than the dendritic subspace error by a constant factor (Fig. S5a-e). For equal somatic and dendritic dimensionalities and loading variances, combining Eqs. S53 and S54 and neglecting the terms involving *g* gives,

,
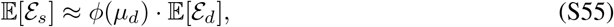

where the constant factor *ϕ*(*µ*_*d*_) can be recognized as the variance of a Bernoulli random variable, which is a function of the baseline burst probability, sig(*µ*_*d*_). This can be understood simply as the approximate information content of a burst per event. This approximation predicts that the dendritic subspace error has the lower bound, E[ℰ_*d*_] 4 · E[ℰ_*s*_], with equality achieved when sig(*µ*_*d*_) = 1*/*2. Figure S5f verifies this dependence on the baseline burst probability and the predicted bound.

### H.5 Sensitivity analysis

As stated in the main text, we used our theoretical results (Eqs. S42 and S52), along with the known data and fitted parameters, to estimate the subspace error regime of the model fits in Fig. 3. Our theory required us to estimate the variance of the true factor loading weights (weight variance) and the dimensionality of each subspace. To do this, we used the factor loading weights from the trained model, 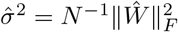, along with its estimated dimensionality 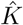 to form a point estimate.

Figure S7a shows the subspace error as a function of these parameters, with all other parameters matched to the VBN data, including the fitted offsets 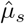 and 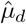 of each neuron. The estimated weight variance and dimensionality are denoted by the grey star. Figure S7b shows the expected subspace error from theory and MC simulations at the point estimate, fitted with both the base and hybrid models. The simulated somatic and dendritic subspace errors were reasonably low, and as expected from theory (Eq. S55), the dendritic subspace error was larger than the somatic subspace error. Similar results were found for the Rodgers data (Fig. S7c-d), but with a lower dendritic subspace error than the VBN data. Similar results were found for the additional recording sessions that were fit (Fig. S4).

It should be noted that our sensitivity analysis makes several simplifying assumptions: we derived subspace error only for the base model, assumed all neurons were burst-coding, and assumed Gaussian-distributed weights and inference errors. Although hybrid-model simulations agreed with the theory and the Gaussian error approximation was appropriate in the low-error regime, these assumptions may lead to optimistic estimates of sensitivity.

## I Model extensions

Although MFA provides a biologically motivated and interpretable framework for investigating burst codes, our current models make several simplifying assumptions. Existing latent variable methods for analyzing firing rates provide natural ways to extend our framework to incorporate shared inputs, temporal correlations, and spike-history effects. Here, we outline three promising directions for future work.

### I.1 A model with shared and private factors

The success of the hybrid model (Fig. 3), together with evidence that multiple pathways target both somatic and dendritic compartments, motivates a model in which event rates and burst probabilities depend on both shared and private factors. Given the emission model from Eq. 2, we propose a model where the event rate and burst probability are given by,

,
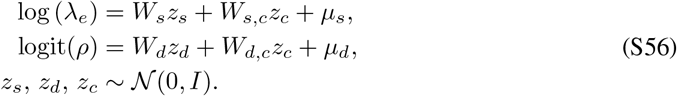

Where 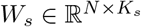 and 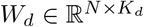 are the private factor loading weights, and 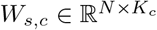 and 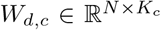 are the shared factor loading weights (common loading weights). Similarly, 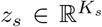 and 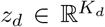 are the private factors, and 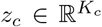 are the shared factors (common factors). As before, *µ*_*s*_, *µ*_*d*_ ∈ ℝ^*N*^ define the means, and nonlinear functions are applied point-wise to map each linear-Gaussian model onto the correct domain.

### I.2 A dynamic latent variable model

Our current implementation of MFA uses 250 ms time bins and assumes temporally independent latent factors. Although this is sufficient for estimating population subspaces, modelling temporal correlations would allow us to investigate population trajectories and their timescales. We therefore propose extending MFA with dynamic latent factors, drawing on existing latent dynamical models [11, 14, 42, 45, 46].

The simplest extension would replace the independent Gaussian priors with Gaussian process (GP) priors, as in Gaussian Process Factor Analysis (GPFA) [14, 45, 46]. For Gaussian Process MFA (GPMFA), we consider each latent dimension to follow

,
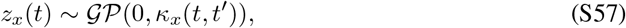

where *κ* is the covariance kernel, typically given by the squared exponential,

,
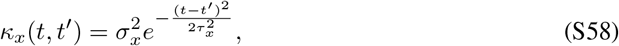

Where 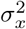 is variance, *τ*_*x*_ is the smoothness timescale, and *x* ∈ {*s, d, c*}.

### I.3 A continuous-time marked point-process model for burst generation

The emission model in Eq. 2 can be viewed as a coarse-grained marked point process. We can therefore extend MFA to continuous time by modelling event onsets as an inhomogeneous Poisson process with intensity *λ*_*e*_(*t*), where each event is marked as a burst with probability *ρ*(*t*). Specifically,

,
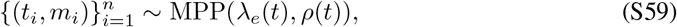

where *t*_*i*_ denotes the event times, *n* is the total number of events, and *m*_*i*_ ∈ {0, 1} is a binary mark indicating whether an event is a burst, with *m*_*i*_ | *t*_*i*_ ~ Bernoulli(*ρ*(*t*_*i*_)). The time-varying event rate and burst probability could then be parameterized using the shared and private factors and Gaussian process priors in Eqs. S56 and S57.

Alternatively, we could introduce a history-dependent marked point process with an explicit mechanism for generating intra-burst spikes, such as the bursting spike-response model [41]. This approach would allow burst identities and lengths to be inferred directly from spike times, without requiring prior burst classification. Furthermore, because the model would account for all spikes, it could be compared directly with competing firing-rate point-process models.

## Footnotes

1 An inhomogeneous Poisson process can produce bursts purely by chance, with a burst fraction proportional to the underlying firing rate.

2 The event and burst distinction used here is still a useful decomposition of spike times, even if bursts are generated from a different mechanism. However, the emission model should be modified to account for this difference.

## Notes

### Competing Interest Statement

The authors have declared no competing interest.

### Summary of Updates

We added a hybrid model that allows burst probability to depend on both event rate and separate latent factors. We expanded the simulations to test how well the model detects burst coding when only some neurons in a population use it. We reanalyzed the mouse visual change detection recordings and added an analysis of recordings from mice performing a whisker-based shape discrimination task. In both datasets, the hybrid model predicted bursting better than the null model, although the improvement varied across sessions. We also updated the sensitivity analysis and expanded the discussion of limitations.

